# SUMO-mediated regulation of a H3K4me3 reader controls germline development in *C. elegans*

**DOI:** 10.1101/2024.02.27.582283

**Authors:** Cátia A. Carvalho, Ulrike Bening Abu-Shach, Asha Raju, Zlata Vershinin, Dan Levy, Mike Boxem, Limor Broday

## Abstract

ULP-2 is a conserved SUMO protease required for embryonic development in *C. elegans*. Here we revealed that ULP-2 controls germline development by regulating the PHD-SET domain protein, SET-26. Specifically, the *ulp-2* mutant hermaphrodites exhibit increased sterility and progressive elevation in global protein sumoylation. In the progeny of homozygous animals, meiosis is arrested at the diplotene stage and the cells in the proximal germline acquire a somatic fate. Germline RNAseq analysis revealed the downregulation of numerous germline genes, whereas somatic gene expression is upregulated in *ulp-2* mutant gonads. To determine the key factors that are regulated by ULP-2, we performed a yeast two-hybrid screen and identified the H3K4me3 reader, SET-26. Genetic interaction was observed in double mutant *ulp-2*;*set-26* resulting in enhanced sterility phenotype to complete sterility in the first generation of homozygous offspring. Consistently, SET-26 is sumoylated and its sumoylation levels are regulated by ULP-2. Moreover, we detected reduction in H3K4me3 levels bound to SET-26 in the *ulp-2* mutant background. A comparative proteomics screen between WT and *ulp-2* loss of activity identified the predicted methyltransferase SET-27 as a ULP-2-dependent SET-26-associated protein. SET-27 knockout genetically interacts with ULP-2 in the germline, but not with SET-26. Taken together, we revealed a ULP-2/SET-26 axis which is required for the maintenance and regulation of germline development.

## Introduction

SUMO, a small ubiquitin-like modifier, is covalently attached to target proteins [1,2] and is essential for development and viability in eukaryotes [3]. SUMO modification is highly dynamic and reversible; its deconjugation is mediated by specific SUMO proteases that cleave the isopeptide bond between the SUMO moiety and substrates [4,5]. The important role of SUMO in the regulation of gene expression has been extensively demonstrated [6], and many transcription factors and chromatin regulators have been identified as SUMO target proteins [7] [8]. Sumoylation is associated with both transcriptional repression and activation [9]. Post-translational histone modifications, including methylation, are epigenetic marks that generate binding sites for reader proteins, which transduce these signals to downstream effects [10,11]. Studies on histone H3 methylation revealed that methylation on lysine residues K4, K36, and K79 are often linked to transcriptional activation, whereas methylation events on histone H3 on lysine residues K9, K20, and K27 are associated with gene repression [11]. It has been demonstrated that the SUMO machinery contributes to the regulation of repressive histone methylation; for example, sumoylation has been shown to be required for Polycomb group protein (PcG) Pc2 recruitment to H3K27me3 marks in mouse embryonic fibroblast cells [12] and for the activity of the PcG-like protein SOP-2 in *C. elegans* [13]. Pc2 has been shown to act as a SUMO E3 ligase [14]. Downregulation of sumoylation by Ubc9 knockdown in embryonic stem cells leads to genome-wide loss of H3K9me3-dependent heterochromatin [15]. In *Drosophila,* deposition of H3K9me3 marks depends on SUMO and the PIAS SUMO E3 ligase Su(var)2–10, which recruits the SetDB1/Wde complex [16]. Many reader proteins of histone methylation have been identified [17]; among them is the chromodomain (CD), found in PcG proteins and the heterochromatin protein 1 (HP1); the CD plays an important role in maintaining repressed chromatin states [18,19]. In *C. elegans,* knockdown of SUMO alters the chromatin-binding pattern of the chromodomain protein MRG-1 [20]. An additional histone methylation reader family is the plant homeodomain (PHD) zinc fingers; this includes ∼100 distinct PHD fingers with histone-binding activity [21]. Structural studies revealed that a modified histone H3 tail is extensively associated with the PHD finger domain, providing a high degree of specificity of this domain to the active mark H3K4me3 [22,23]. PHD domains were suggested to function as SUMO E3-ligases [24]. Several histone readers contain more than one “reading-module” and are therefore potentially able to receive and translate complex signals. One of them is the mixed lineage leukemia 5 protein (MLL5), which contains both a Su(var)3–9, Enhancer-of-zeste, Trithorax (SET) domain, and a PHD domain [25]. MLL5 binds to H3K4me3 marks through the PHD domain and associates with chromatin downstream of the transcriptional start sites of active genes [26-28]. Based on a similar sequence of their SET domains, MLL5 appears to be the mammalian homolog of yeast SET3/SET4 paralogs, *Drosophila* UpSET, and the *C. elegans* SET-9/SET-26 paralogs [29,30]. The unique property of the SET domain in these proteins is the lack of histone methyl transferase activity [25,31,32](and this study). The SET domains of SET3 and UpSET were shown to be associated with histone deacetylases [31,32] and to regulate H3K9me2 levels [33].

*C. elegans* SET-9 and SET-26 are two histone readers with a PHD-SET domain that binds to H3K4me3 marks in germline and somatic genes [30]. Even though they are 97% identical in their sequence, they perform different functions. Both SET-9 and SET-26 synergize to maintain germline function across generations; however, SET-26 plays an additional role in regulating lifespan in a germline-independent manner and in regulating resistance to heat stress [30,34,35]. The *C. elegans* ULP-2 protein occupies the Ulp2-like branch of the SUMO proteases and shares 17% similarity with the catalytic domains of the mammalian SUMO proteases SENP6 and SENP7 [36]. Proteomics studies revealed that SENP6 is associated with chromatin organization complexes [37] and is required to stabilize the centromeric H3 variant CENP-A [38,39]. Here we revealed a role for the ULP-2 SUMO protease in regulating the active H3K4me3 histone methylation mark in the germline. Importantly, we demonstrated that ULP-2 plays a key role in germline development and maintenance of germ cell fate; we identified the SET-26 histone reader as a central target for this activity.

## Results

### Increased sumoylation levels in homozygous *ulp-2* mutant animals inversely shapes fertility output and lifespan

We previously reported that the homozygous progeny of the heterozygous hermaphrodites of a mutant allele of ULP-2 SUMO protease, *ulp-2(tv380)*, a predicted null allele, is arrested during embryonic epidermal morphogenesis with incomplete penetrance [36]. We observed that the escaper group that completed embryonic epidermal morphogenesis (∼37%, Figure 1A) continued to develop to sterile adults probably due to a sufficient maternal product. The entire developmental process was variable between the progeny of homozygous animals and was associated with a developmental delay (8-9 days, Figure 1A). Since we observed sterility in the second homozygous generation (F2 *ulp-2(tv380)*), we measured the brood size (unhatched embryos and larvae) of *ulp-2* mutant animals. We observed that the first homozygous generation of *ulp-2* mutants (F1 *ulp-2(tv380)*) had already decreased its fertility (∼55% reduction in fertility, 153±53 progeny) compared with WT animals (277±30 progeny), whereas the second generation of homozygous mutants (F2 *ulp-2(tv380)*) was almost completely sterile (∼99% sterility,0.2±2 progeny) (Figure 1B). This suggests that there is a reproductive decline in the homozygous generations of *ulp-2* mutant animals.

**Figure 1.**
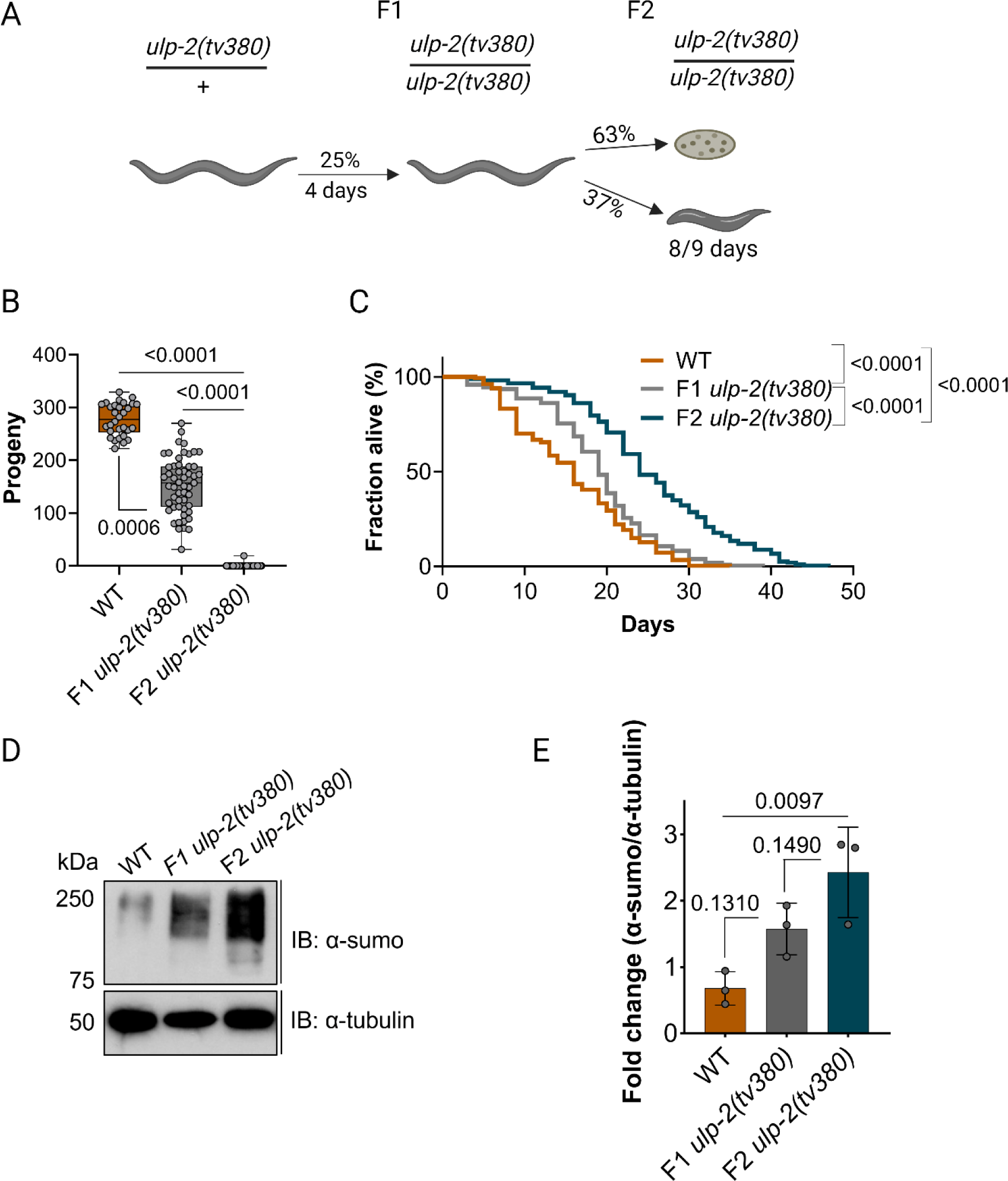
Loss of function of ULP-2 induces fertility decline and increases longevity accompanied by the accumulation of SUMO conjugates. **A.** Schematic representation of the developmental progress of *ulp-2(tv380)* animals. B. Quantification of the number of progeny laid by WT (30 animals), F1 *ulp-2(tv380)* (53 animals), and F2 *ulp-2(tv380)* (94 animals) in 2 biological replicates; the Shapiro-Wilk and one-way ANOVA tests on ranks (Kruskal-Wallis), followed by Dunn’s post-hoc test, were used; ns=p>0.05. C. Survival curves for WT (252 death events+100 censored), F1 *ulp-2(tv380)* (220 death events+162 censored) and F2 *ulp-2(tv380)* (213 death events+228 censored) animals in 2 biological replicates; the log-rank (Mantel-Cox) test was used; ns=p>0.05. D. Representative immunoblot of the total sumoylation levels of WT, F1 *ulp-2(tv380),* and F2 *ulp-2(tv380)* animals; 3 biological replicates; the upper panel was probed with anti-sumo and the bottom panel was probed with anti-α-tubulin. E. Quantification of the total levels of sumoylated proteins in D.; the Shapiro-Wilk and one-way ANOVA tests, followed by Tukey’s post hoc statistical test, were used; ns=p>0.05.

Since the reproductive system plays a key role in regulating the lifespan of the organism [40,41], we determined whether the lifespan of the *ulp-2* mutants was affected. We found that the median lifespan of WT animals is 16 days, compared with 19 days in F1 *ulp-2(tv380)* and 24 days in F2 *ulp-2(tv380)* animals (Figure 1C, Table S1). These results reflect a progressive increase in the lifespan of animals lacking ULP-2 activity, which correlates inversely with its reproductive function. Since both generations of homozygous animals are genetically identical, we reasoned that in addition to the reduced maternal product, the phenotypic differences could have arisen from an accumulation of SUMO conjugates over time. Indeed, whereas F1 *ulp-2(tv380)* mutant animals contained double the amount of SUMOylated substrates (mean=1.6±0.4) compared with WT (mean=0.7±0.3); the F2 *ulp-2(tv380)* mutant animals contained triple the amount of SUMO conjugates (mean=2.4±0.7) (Figure 1D-E). All together, these results suggest that the accumulation of SUMO conjugates in *ulp-2* mutant animals contributes to the decline in fertility concomitantly with an increase in lifespan.

### Loss of ULP-2 activity disrupts the transcriptional program in the germline

To characterize the sterility phenotype of *ulp-2* mutant animals, we began by performing transcriptomics analysis of isolated germlines. Morphologically, the germlines of F2 *ulp-2(tv380)* animals have a reduction of ∼50% in length (269 ± 58 µm) compared with WT germlines (533 ± 73 µm) (Figure S1A). The mutant germlines also harbor 45% fewer nuclei, indicating that ULP-2 loss of function negatively affects the cellular content of the germline (Figure S1B). The transcriptomics analysis revealed that a total of 7,202 genes are differentially expressed in the germlines of F2 *ulp-2(tv380)* relative to WT, with 3,577 genes downregulated and 3,625 genes upregulated (Figure 2A, Figure S1C, and Tables S2-S3). Gene Ontology (GO) analysis revealed that the downregulated genes in F2 *ulp-2(tv380)* germlines are associated with biological processes essential for germline maintenance, such as cell cycle regulation, P-granule assembly as well as oocyte development and specification (Figure S2A). On the other hand, the GO analysis of the genes upregulated in the F2 *ulp-2(tv380)* germlines are associated with somatic processes such as immune response, cuticle development, and neurogenesis (Figure S2B). The intersection of our transcriptomics dataset with two previously defined datasets [42], substantiates the GO analyses; whereas the downregulated genes in the F2 *ulp-2(tv380)* germlines largely overlap with the “germline-enriched genes” dataset (R.F.=2.6), the upregulated genes significantly intersect with the “somatic-specific genes” dataset (R.F.=2.4) (Figure 2B). Taken together, these observations suggest that the accumulation of SUMO conjugates in *ulp-2* mutant germlines induces an overall loss of transcripts required to maintain a germline coupled with a permissive somatic transcription.

**Figure 2.**
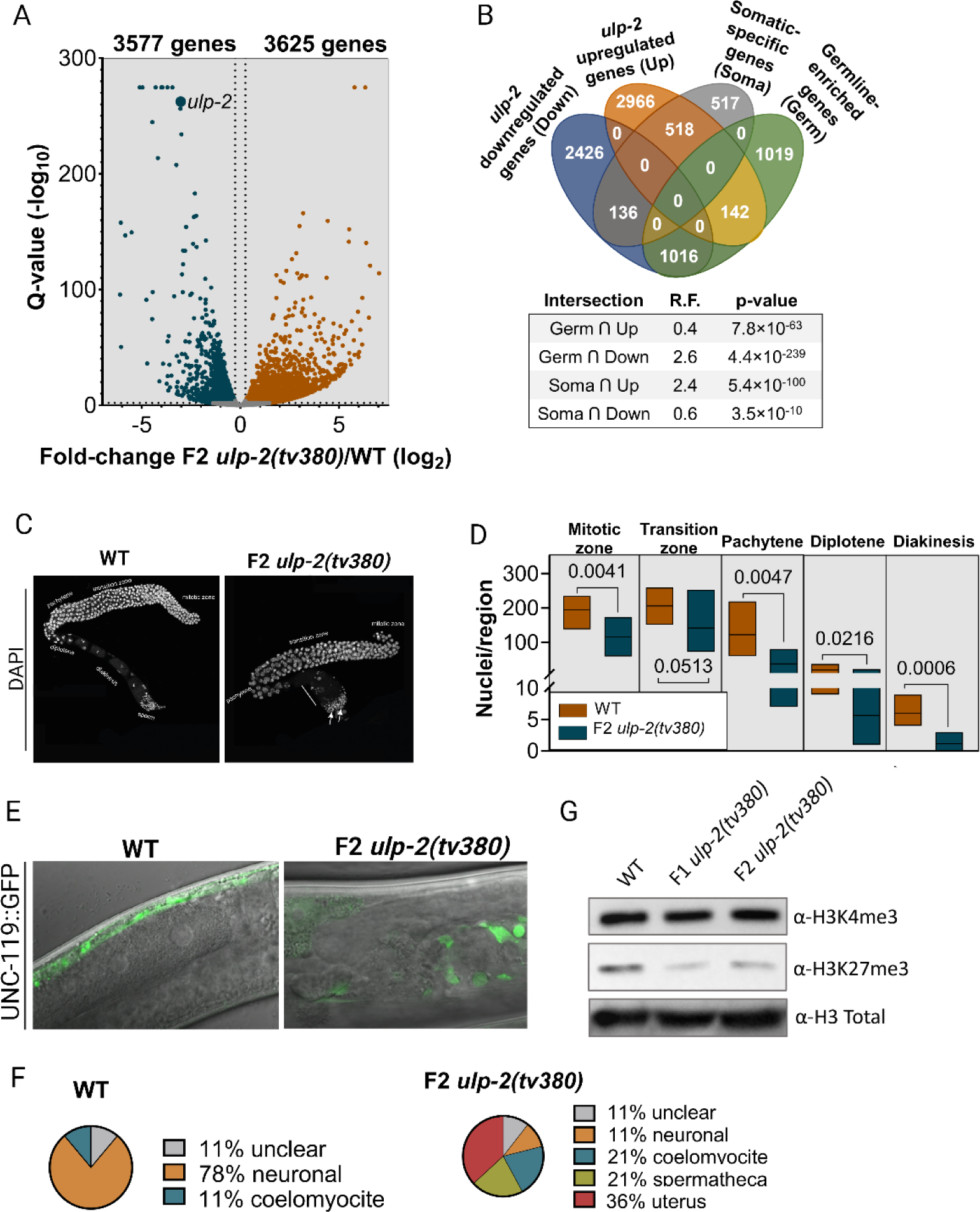
Loss of ULP-2 disrupts the germline gene expression program, leading to impaired oocyte formation and loss of germ cell identity. **A.** A volcano plot displaying the distribution of the differentially expressed genes of the F2 *ulp-2(tv380)* germlines in comparison to WT. The cut-off values used were FDR>2 (-log10) and fold change>|1.2| (log2). **B.** A Venn diagram representing the intersection between downregulated and upregulated genes in the F2 *ulp-2(tv380)* germlines and the established datasets of germline-enriched and somatic-specific genes [42]; the representation factor (R.F.) of each intersection and the associated p-value are shown; Fisher’s exact test was used. **C.** Representative images of the isolated germlines of WT and F2 *ulp-2(tv380)* animals; the germline zones are indicated; abnormal sperm and large nuclei in the proximal gonad are denoted by white arrows in the *ulp-2(tv380)* germline. Scale bar=100µm. **D.** Quantification of the number of nuclei per region in the germlines of WT (7 germlines, 3,834 nuclei) and F2 *ulp-2(tv380)* (6 germlines, 1,809 nuclei) animals; the Shapiro-Wilk and the two-tailed Mann-Whitney U tests were used; ns=p>0.05. **E.** Representative images of UNC-119::GFP localization in WT and F2 *ulp-2(tv380)* animals. **F.** Distribution of UNC-119::GFP localization in WT (8 worms) and F2 *ulp-2(tv380)* (13 worms) animals from **E**. **G.** Representative immunoblot of the total amount of H3K4me3, H3K27me3, and Histone 3 in WT, F1 *ulp-2(tv380),* and F2 *ulp-2(tv380)* animals; 3 biological replicates.

### Impaired desumoylation prevents gamete formation and the maintenance of germ cell identity

To dissect the major germline defects we analyzed DAPI-stained germlines. The germline of F2 *ulp-2(tv380)* proliferate, but they are relatively smaller and gametogenesis is arrested (Figure 2C). The mutant germlines appear to be relatively healthy at the L4 stage but deteriorate, along with high variability when adults. The number of germ cells undergoing mitosis was reduced by 41% in F2 *ulp-2(tv380)* animals, indicating that fewer cells undergo meiotic differentiation. Accordingly, we observed a sharp reduction in the number of nuclei in all meiotic stages. The most dramatic reduction was in diakinesis, with 81% fewer nuclei, resulting in an almost complete absence of oocytes formed in F2 *ulp-2(tv380)* germlines (Figure 2C-D). In *C. elegans*, when fertilization is abnormal or there is a premature depletion of sperm, it is possible to detect unfertilized oocytes that were activated spontaneously and that underwent multiple DNA replication rounds [43]. However, we could not detect such endomitotic oocytes in the *ulp-2(tv380)* gonads, suggesting that the blockage occurred before oocyte meiotic maturation and fertilization, leading to complete sterility. In addition to sperm cells that appear round with a uniform compact size, we identified apparently undifferentiated larger misshapen nuclei appearing in variable sizes and numbers in the proximal gonad (Figure 2C, arrows), which resembles the *lin-41* mutant phenotype [44]. To determine whether these larger cells lost their germline fate, we crossed the *ulp-2* mutant animals with the pan-neuronal marker UNC-119::GFP [45], which has been used to detect germ cell reprogramming to a neuron-like somatic fate [46-48]. We observed germline expression of UNC-119::GFP in these cell clusters with axonal-like cellular extensions (Figure 2E-F) that were detected only in the proximal gonad, suggesting that the role of ULP-2 in protecting germ cell fate is essential in the proximal germline.

De-repression and ectopic transcription of somatic genes in the germline is a phenomenon that has been observed in the context of impaired chromatin regulation and loss of germline fate [49]. To examine possible alterations in chromatin marks, we measured the levels of the histone marks broadly accepted as transcriptionally permissive and repressive, H3K4me3 and H3K27me3, respectively [50,51]. Although the H3K4me3 levels were unaffected in both generations of *ulp-2(tv380)* homozygous animals, the levels of H3K27me3 were reduced compared with WT (Figure 2G). Therefore, the induction of somatic gene transcription in *ulp-2* mutant germlines may originate from this global loss of H3K27me3 as previously shown for this histone mark [52].

### ULP-2 interacts with the SET-26 H3K4me3 reader to maintain germline function

To identify targets for the SUMO protease activity of ULP-2 in the germline, we performed a yeast two-hybrid (Y2H) screen of a *C. elegans* mixed-stage cDNA library using the N-terminal domain of ULP-2 as bait. We identified SET-26 and a homolog protein, Y73B3A.1. SET-26 is a PHD-SET protein and Y73B3A.1 has a high sequence homology to SET-26 but lacks the PHD-SET domain (Figure S3A). SET-9 and SET-26 are paralogs that have been established as H3K4me3 readers required for a functional germline [30]; therefore, they would be likely targets for ULP-2 activity in the germline. To examine the specificity of ULP-2 interaction with SET-26 and Y73B3A.1, first, we simultaneously downregulated SET-9, SET-26, and Y73B3A.1 (“Set3” RNAi) in homozygous F1 *ulp-2(tv380)* and observed a sharp decrease in the average number of progeny produced and ∼40% sterility when compared with F1 *ulp-2(tv380)* and WT (Figure S3B-C). Conversely, *ulp-2* downregulation by RNAI in individual mutants of each family member led to synergistically increased embryonic lethality and near sterility between ULP-2 and SET-26, but not with SET-9 and Y73B3A.1 (Figure S3D-E). We therefore decided to focus on the interaction of ULP-2 with SET-26. To confirm the interaction of SET-26 with ULP-2 and map the binding region, we tested the N-terminal (exons 1–4) and the C-terminal (exons 6–9) regions of SET-26 for interaction with ULP-2 by Y2H (Figure S3A). The results indicated that the N-terminal region of SET-26 mediates its interaction with the N-terminal domain of ULP-2 (Figure 3A). To further examine the genetic interaction between ULP-2 and SET-26, we generated double mutants of *ulp-2(tv380)* with two loss of function alleles of *set-26*. In these two alleles, the N-terminal is intact and the deletion causes early termination of the reading frame before or inside the PHD domain, leading to a shorter protein with no PHD and SET domains (Figure S3F). Double mutants *ulp-2;set-26* rendered the first generation of homozygous *ulp-2(tv380)* near sterile, resembling the F2 *ulp-2(tv380)* phenotype (Figure 3B). F1 homozygous *ulp-2(tv380)* animals express sufficient maternal ULP-2 for germline development, which is abolished in the background of the *set-26* loss of function alleles. These results suggest that SET-26 is a key target of ULP-2 activity in the germline. In addition to its function in the germline, SET-26 has been shown to regulate somatic-driven maintenance of a normal lifespan independently of the germline [34,35]. To determine whether ULP-2 and SET-26 interaction extends to somatic tissues, we measured the lifespan of the double mutants. However, we did not observe a synergy between ULP-2 and SET-26 in extending lifespan (Figure 3C, Table S1). Overall, this suggests that the synergistic interaction between SET-26 and ULP-2 occurs primarily in the germline and is required for its maintenance.

**Figure 3.**
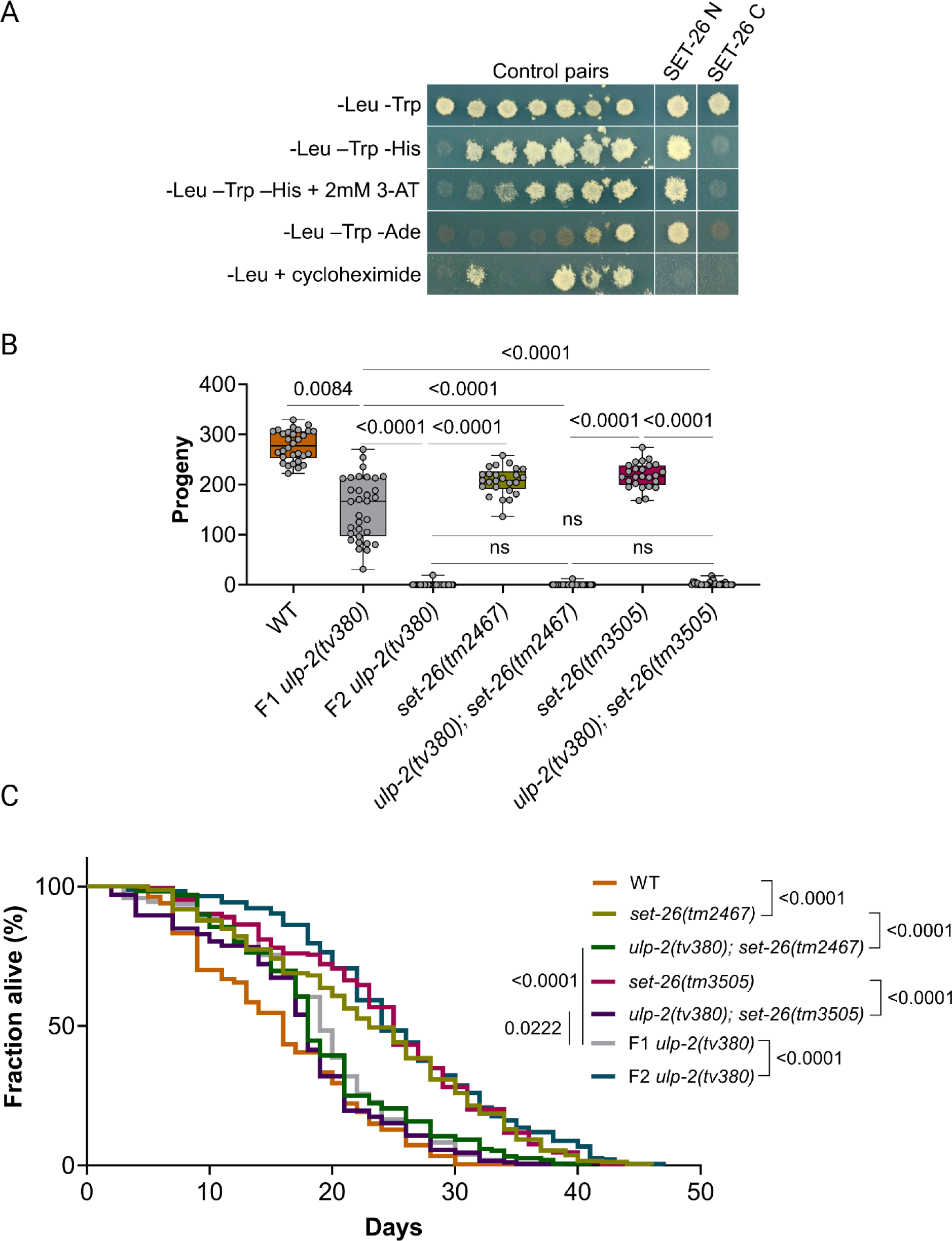
SET-26 is required for maintenance of the germline function of ULP-2. **A.** A yeast two-hybrid analysis of the interaction between the N- and C-terminal domains of SET-26 and the N-terminal domain of ULP-2. The top row is a non-selective growth plate. The middle three rows are selective plates that only allow yeast growth in the presence of an interaction. The bottom row is a control plate where growth indicates the auto-activation of reporter genes by the DNA-binding domain protein fusion. Controls are the protein pairs of a known reporter activation strength. **B.** Quantification of the number of progeny produced by WT (30 animals, 277±30 offspring per mother), *ulp-2(tv380)* (32 animals, 153±63 offspring per mother), F2 *ulp-2(tv380)* (94 animals, 0.2±2 offspring per mother), *set-26(tm2467)* (25 animals, 207±27 offspring per mother), *ulp-2(tv380);set-26(tm2467)* (39 animals, 0.3±2 offspring per mother), *set-26(tm3505)* (25 animals, 218±25 offspring per mother) and *ulp-2(tv380);set-26(tm3505)* (37 animals, 2±4 offspring per mother); 2 biological replicates; the Shapiro-Wilk and one-way ANOVA tests on ranks (Kruskal-Wallis), followed by Dunn’s post-hoc test, were used; ns=p>0.05. **C.** Survival curves for WT (252 death events+100 censored animals), *ulp-2(tv380)* (220 death events+162 censored animals), F2 *ulp-2(tv380)* (213 death events+228 censored animals), *set-26(tm2467)* (254 death events+92 censored animals), *ulp-2(tv380);set-26(tm2467)* (157 death events+210 censored animals), *set-26(tm3505)* (245 death events+81 censored animals), and *ulp-2(tv380);set-26(tm3505)* (205 death events+188 censored animals); the log-rank (Mantel-Cox) test was used; ns=p>0.05.

### ULP-2 controls the sumoylation levels of SET-26

SUMO proteases mediate the enzymatic removal of SUMO moieties from sumoylated proteins [53]. The physical and genetic interaction between SET-26 and ULP-2 suggests that SET-26 is a target of ULP-2 protease activity. To investigate this hypothesis, we first assessed whether SET-26 is a SUMO-acceptor protein. Using an in-vitro sumoylation assay, the SET domain with or without the adjacent PHD domain of SET-26 accumulated multiple higher molecular weight bands detected by anti-SUMO antibody, suggesting that SET-26 can be either sumoylated on several lysine residues or it can be polysumoylated (Figure 4A). The reaction proceeded more rapidly when the region including the PHD-SET domain of SET-26 was subjected to the sumoylation reaction (Figure 4A), suggesting that the PHD domain in SET-26 may act as an intramolecular SUMO E3 ligase, as previously shown for the sumoylation of the bromodomain of the KAP1 corepressor [24] to enhance the sumoylation of the SET domain. Next, we performed in-vitro desumoylation reactions and found that incubation of sumoylated PHD-SET domains with ULP-2 resulted in the removal of the SUMO peptides, as manifested by a sharp decrease in signal, compared with the sumoylated input (Figure 4B). Overall, these results indicate that ULP-2 can regulate the sumoylation levels of SET-26 in-vitro. If SET-26 is a target of ULP-2 in-vivo, increased levels of sumoylated SET-26 should be observed in the *ulp-2* loss of function background. Immunoprecipitation of SET-26::GFP did not exhibit strongly reacting bands with anti-SUMO antibody in a WT background (Figure 4C). On the other hand, SET-26::GFP immunoprecipitated from F2 *ulp-2(tv380)* mutant animals exhibited a strong reactivity with the anti-SUMO antibody, corresponding to a 3-fold increase in sumoylation levels (Figure 4C-D). These results suggest that ULP-2 regulates the sumoylation levels of SET-26 in *C. elegans*. SET-26 is the ortholog of human MLL5, fly UpSET, as well as yeast SET3 and SET4 that lack histone methyltransferase activity [25,31,32]. We determined whether the sumoylation of the PHD-SET domain of SET-26 can activate the SET methyltransferase in-vitro, but no such activity was detected (Figure S4).

**Figure 4.**
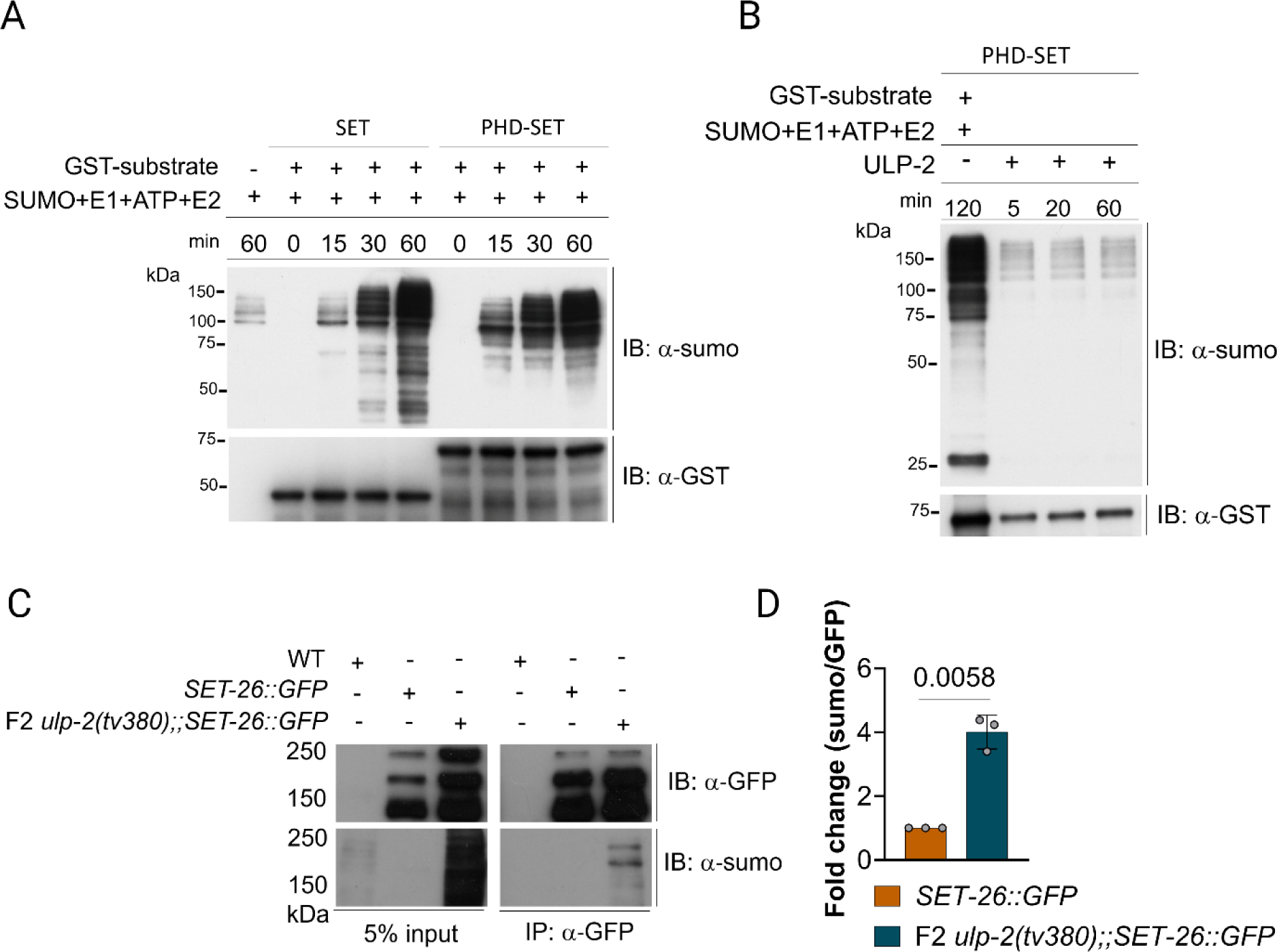
SET-26 is a SUMO target and is desumoylated by ULP-2. **A.** In-vitro sumoylation of the SET and PHD-SET domains of SET-26. Bacterially expressed GST-SET and GST-PHD-SET were incubated with E1 (SAE1/SAE2), E2 (UBC9), SUMO1, and ATP for 0, 15, 30, and 60 min at 37°C. Control reactions were incubated for 60 min without the substrate. The anti-SUMO and anti-GST antibodies were used to detect sumoylation and the GST proteins, respectively. **B.** The ULP-2 catalytic domain desumoylates the PHD-SET domain in vitro. The GST-PHD-SET was sumoylated for 2 hr before adding ULP-2 for the indicated time points. Reactions were carried out at 37°C. The anti-SUMO and anti-GST antibodies were used similarly to A. **C.** SET-26 in-vivo sumoylation in WT and F2 *ulp-2(tv380)* backgrounds; anti-GFP and anti-SUMO antibodies were used to probe SET-26::GFP and the sumoylation levels of SET-26, respectively. **D.** Quantification of the normalized levels of the sumoylation of SET-26 in **C.** for 3 biological replicates; one sample t-test was used; ns=p>0.05.

### SET-26 binding to H3K4me3 marks depends on ULP-2

In the nucleus, SET-26 binds to H3K4me3 marks to regulate gene expression [30]. Hence, we intersected a CHIP-seq dataset of the SET-9 and SET-26 genomic binding sites (85% SET-26 binding sites) [30] with our transcriptomics dataset. Interestingly, although we did not observe a meaningful intersection with the upregulated genes (R.F.=8), the overlap between the pool of downregulated transcripts in the F2 *ulp2(tv380)* mutant germlines and the genes bound by SET-9/26 was significantly overrepresented (R.F.=2.2) (Figure S5A). This suggests that SET-26 can influence the gene expression program in the germline that we found to be downregulated in the F2 *ulp2(tv380)* mutant germlines. Thus, we hypothesized that if the SET-26 reader function is altered by an excess of its sumoylation, it can mediate the disruption of the germline gene expression program, leading to the sterility of the F2 *ulp2(tv380)* animals. To test this hypothesis, we evaluated SET-26 de facto reader ability by measuring its affinity for H3K4me3 marks. Consistent with a previous report [30], immunoprecipitated SET-26::GFP was found to be bound to the H3K4me3 marks in WT (Figure 5A). However, in F2 *ulp-2(tv380)* mutant animals, SET-26::GFP binding to H3K4me3 was, on average, reduced by 55%, compared with WT (Figure 5A-B). Taking into consideration that ULP-2 activity does not alter the global levels of H3K4me3 (Figure 2G), these results indicate that excessive sumoylation of SET-26 decreases its H3K4me3 binding affinity, which potentially mediates the disruption of the germline gene expression program observed upon ULP-2 loss of function.

**Figure 5.**
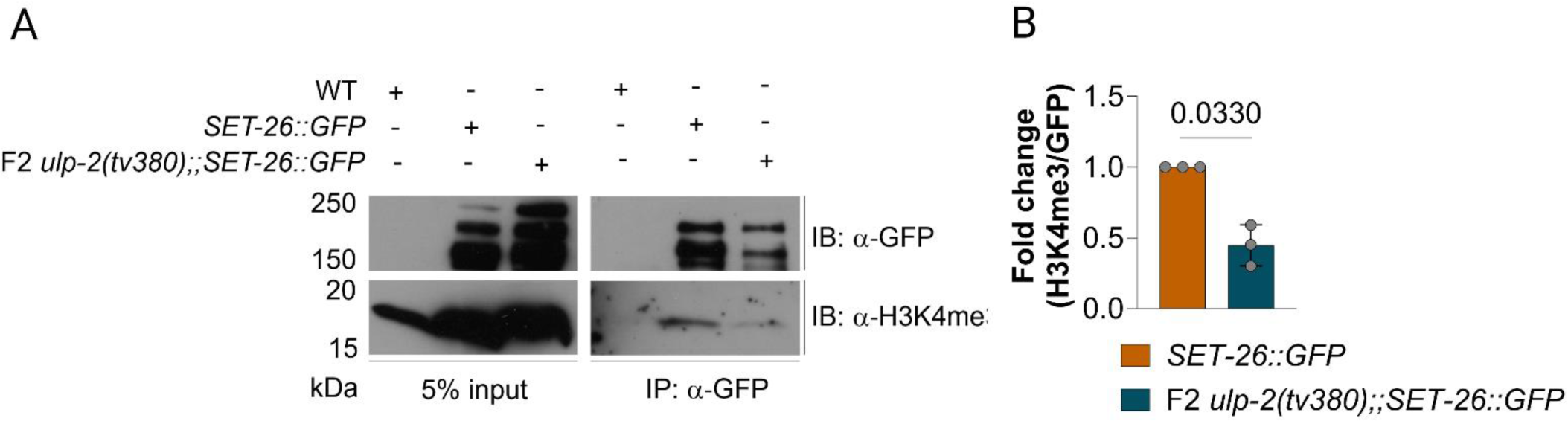
The sumoylation levels of SET-26 regulate its H3K4me3 reader capacity. **A.** Immunoblot of SET-26::GFP binding to H3K4me3 marks in WT versus F2 *ulp-2tv(380)* backgrounds; anti-GFP and anti-H3K4me3 antibodies were used to recognize the levels of SET-26::GFP and H3K4me3 bound to SET-26::GFP, respectively. **B.** Quantification of the normalized levels of H3K4me3 binding to SET-26 in **A.**; 3 biological replicates; One-sample t-test was used; ns=p>0.05.

### Excessive sumoylation of SET-26 weakens its protein-protein interactions

Sumoylation can alter the function of a given protein in multiple ways; for instance, it can alter its subcellular localization, stability, or its interacting partners [9]. Besides its PhD and SET domains, SET-26 is predicted to be mostly composed of disordered protein regions. These regions assemble multiple protein-protein interactions and are also favored protein regions for SUMO conjugation events [54,55]. To determine whether excessive SET-26 sumoylation could alter its protein interactions, we compared the interactome of SET-26::GFP in the condition of *ulp-2(RNAi)* versus WT through immunoprecipitation and mass spectrometry (Figure 6A, Figure S5B-C). In WT, SET-26 was found to be in a complex with several proteins (Figure 6B, right side) whose general biological function concerns metabolic regulation (Figure S5D). However, when SET-26 is excessively sumoylated (*ulp-2(RNAi*), Figure 6A), we observed a broad decrease in its ability to bind to its WT interacting proteins, and there was only a slight gain in new interactors (Figure 6B, left side; Table S4). This indicates that excessive sumoylation of SET-26 decreases its complex-forming capacity, which could directly alter SET-26’s ability to either bind to or remain bound to H3K4me3 marks. We chose SET-27 for further analysis due to its predicted methyltransferase activity (https://wormbase.org/). SET-27 showed no obvious phenotype in an RNAi screen of SET domain proteins [56]. We generated a CRISPR/Cas9-mediated knockout of SET-27 (*set-27(tv381)*) to mimic the decreased SET-27/SET-26 interaction observed in the condition of excessive sumoylation (Figure S5E). We observed that when SET-27 function is lost, there is a small increase of ∼7% in the number of progeny produced relative to WT, indicating a minor function of SET-27 in the germline (Figure 6C). Double mutant *set-26;set-27* does not impact the number of progeny generated compared with SET-26 loss of function, probably due to the gain of function nature of sumoylated SET-26 which is functionally different from the *set-26* deletion allele. However, we observed that double mutant *ulp-2;set-27* displayed a reduction of ∼17% in progeny produced compared with F1 *ulp-2(tv380)* mutants (Figure 6C), highlighting a genetic interaction between SET-27 and ULP-2. This suggests that through its interaction with ULP-2, SET-27 may impact SET-26 sumoylation levels and modulate its reader function.

**Figure 6.**
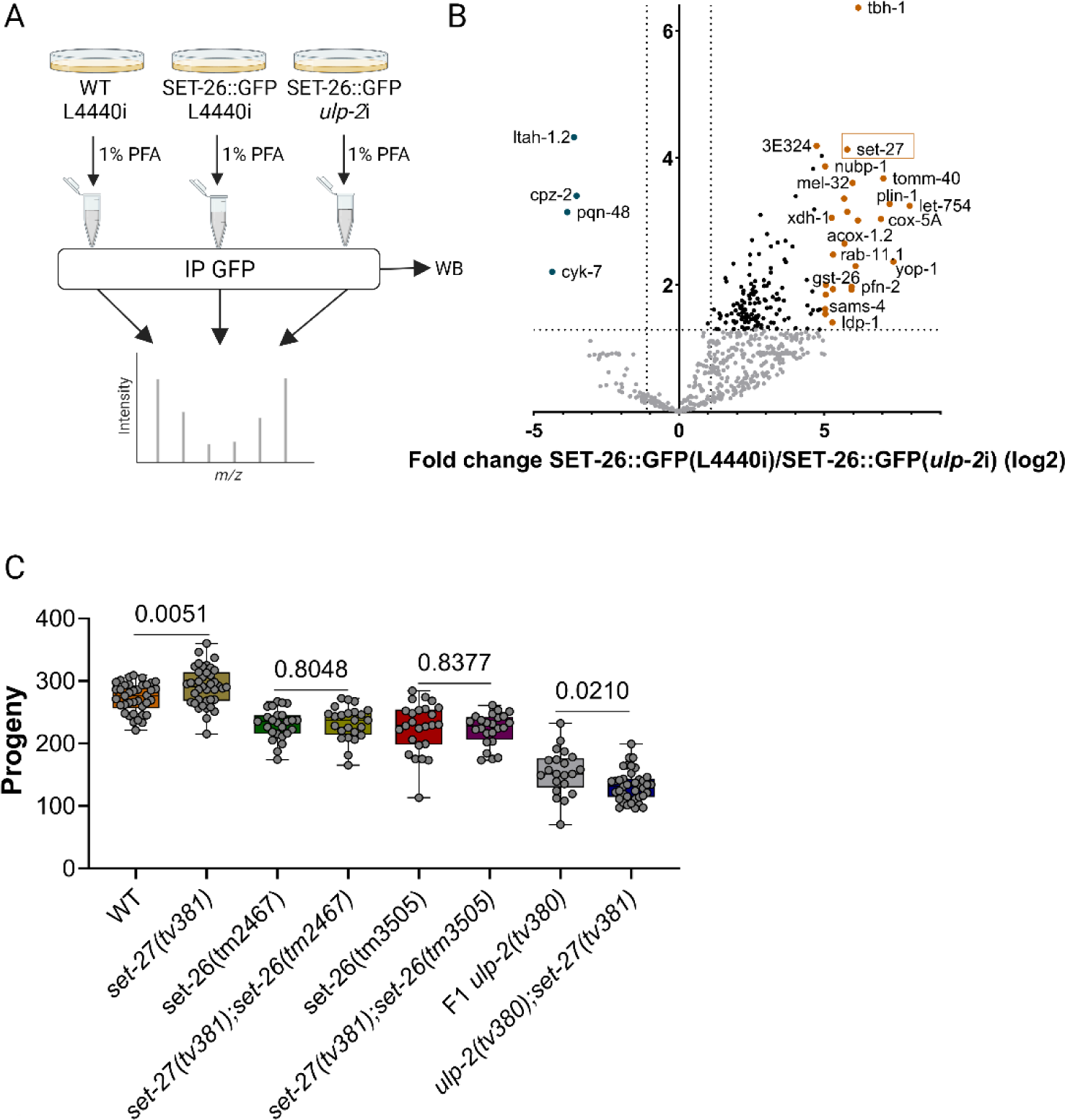
SUMO-dependent disruption of SET-26 complex formation. **A.** Schematic representation of the AP-MS strategy to isolate SET-26::GFP in WT versus an excessive sumoylation background; 3 biological replicates. **B.** A volcano plot depicting the differentially enriched proteins bound to SET-26::GFP in WT versus an excess of SUMOylation backgrounds; the cut-off values used were FDR>1.3 (-log10) and fold change>|1.1| (log2). **C.** Quantification of the number of progeny of WT (38 animals, 273±23 offspring per mother), *set-27(tv381)* (39 animals, 291±31 offspring per mother), *set-26(tm2467)* (25 animals, 230±24 offspring per mother), *set-27(tv381);set-26(tm2467)* (24 animals, 232±27 offspring per mother), *set-26(tm3505)* (25 animals, 225±40 offspring per mother), *set-27(tv381);set-26(tm3505)* (24 animals, 223±25 offspring per mother), F1 *ulp-2(tv380)* (21 animals, 153±36 offspring per mother), and F1 *ulp-2(tv380);set-27(tv381)* (39 animals, 132±25 offspring per mother); the Shapiro-Wilk and two-tailed Welch’s t-tests were used; ns=p>0.05; 2 biological replicates.

## Discussion

In this study we revealed that the SUMO protease ULP-2 is required for germline development and for SET-26 function in the germline. Whereas the first generation of homozygous *ulp-2(tv380)* mutants showed decreased progeny compared with WT, the second generation was completely sterile. This phenotype was accompanied by increased global sumoylation levels, which could directly result from the non-reversible sumoylation of multiple targets or a secondary stress response effect that led to a proteome-wide sumoylation [9]. The most dramatic germline defect was a block in meiosis I and specifically the transition from the end of prophase I to metaphase. Moreover, at this proximal region of the gonad, we also observed the loss of germ cell fate. This suggests that in processes that were found to depend on sumolyation during mitosis in the germline as well as earlier meiotic stages [57], there is a redundancy with other SUMO proteases in order to maintain a balanced sumoylation that is sufficient for germline development and germ cell fate. However, desumoylation processes during the transition from the end of meiotic prophase I are suggested here to be regulated non-redundantly by ULP-2.

Dramatically, transcriptomics analysis of dissected germlines revealed a large-scale downregulation of germline genes together with upregulation of misexpressed somatic genes. The impaired expression of thousands of genes in the germline suggests that knockout of ULP-2 directly affects the chromatin structure in the germline. In total protein extracts we observed a global decrease in H3K27me3 marks that could contribute to the loss of germ cell fate as previously shown [48] as well as to somatic defects as we have recently demonstrated in the mitochondria [58]. However, the late arrest phenotype in germline development of the F2 homozygous *ulp-2* mutant animals differs from the complete elimination of H3K27me3 marks in *mes-2/3/6* mutant which causes germline degeneration [59,60].

Using a yeast two-hybrid screen, we identified the PHD-SET domain protein, SET-26, as a binding partner of ULP-2. SET-26 is a unique SET domain protein with homology to human MLL5 and *Drosophia* upSET [33,61] lacking methyltransferase activity; it has been suggested to act as an H3K4me3 reader [30]. When we compared the CHIP-seq data of SET-9/SET-26 [30] genomic binding sites with our RNAseq *ulp-2* mutant data, we observed that the vast majority of genes that SET-9/SET-26 bind to are downregulated in *ulp-2(tv380)* mutant animals. Moreover, double mutant *ulp-2;set-26* resulted in a dramatic enhancement of the sterility phenotype to complete sterility in the first generation of homozygous animals, thus supporting a genetic interaction between *ulp-2* and *set-26*. To elucidate the functional interaction between ULP-2 and SET-26, we investigated whether SET-26 is sumoylated. We found that it is sumoylated on several residues or is polysumoylated, since we observed multiple bands on Western blots. In-vivo, we observed an increase in SET-26 sumoylation in the *ulp-2* mutant background. Next, we showed that constitutive sumoylation of SET-26 interfered with the direct binding of SET-26 to the H3K4me3 histone marks and its interactors, among them the SET-27 putative methyltransferase. Our study revealed that sumoylation of SET-26 abolished its binding to H3K4me3, leading to impairment of the germline transcriptional program. As previously reported [30], the *set-26* mutant alleles increase lifespan; however, in double mutant *ulp-2;set-26* the lifespan extension was abolished. This suggests that whereas ULP-2 and SET-26 share the regulation of the same germline genes, the two proteins have different chromatin binding sites or transcriptional activities in the chromosomal regions of somatic genes involved in lifespan regulation (Figure S6).

During germline development and maintenance of totipotency, the histone modifiers and histone readers must be tightly regulated. A balance between different histone modifications appears necessary to maintain germline pluripotency [62-65]. In *C. elegans*, the role of SUMO in the epigenetic regulation of the germline was recently demonstrated by the finding that sumoylation of the CCCH zinc-finger PIE-1 and HDAC1 promotes piRNA-dependent transcription silencing and maintains germline fate [66,67]. Another example is the *C. elegans* protein MRG-1, a sumoylated chromodomain protein whose chromatin binding patterns are modified following a decrease in overall sumoylation [20].

Our data suggest that ULP-2 contributes to the formation of an epigenetic barrier that will prevent germ cell reprogramming toward somatic fates using sumoylation-desumoylation cycles. Dynamic sumoylation is an efficient method to strengthen this regulation. Since SET-26 has been shown to bind to and restrict H3K4me3 spread in germline genes [30], it is possible that cycles of sumoylation and desumoylation mediate this binding.

Several studies have demonstrated the role of PHD domains in recognizing histone methylation [68] and in regulating sumoylation [69]. The PHD domain of the KAP1 corepressor functions as a SUMO-E3 ligase of the adjacent bromodomain[24]. The sumoylation of the bromodomain mediates the direct recruitment of H3K9 HMTase and HDAC activities to promoter regions to silence gene expression [24]. SIZ1 is an *Arabidopsis* SUMO E3 ligase and its PHD domain binds to H3K4me3 [70]. The PHD domain of the SET-26 ortholog, MLL5, has been shown to be recruited to actively transcribe genes; it binds specifically to H3K4me3 marks; however, this binding is inhibited by phosphorylation on neighboring residues on the histone (Thr3 or Thr6) [28]. Similarly, as sumoylation modulates protein interactions [8], it is possible that SUMO conjugation masks the interaction surface between the PHD domain of SET-26 and H3K4me3 to regulate the binding of the histone reader to the histone marks.

Histones are also direct substrates of sumoylation. Serial modifications on histones, including H2B ubiquitylation, H3K4 methylation, histone sumoylation, and histone deacetylation, function together to regulate transcription [71]. Histone sumoylation recruits the Set3 histone deacetylase complex to both protein-coding and noncoding RNA (ncRNA) genes [72]. Several SUMO proteases have been shown to regulate histone-modifying enzyme activities during developmental processes. For example, SENP3 regulates the SET1/MLL complex during osteogenic differentiation [73] and the SETD7 histone methyltransferase during sarcomere assembly [74]. Our study suggests that in addition to the sumoylation of the histones themselves and the histone-modifying enzymes, an additional layer of chromatin regulation is contributed by the balanced sumoylation/desumoylation of the readers of specific histone marks.

## Methods

### *C. elegans* strains and genetics

*C. elegans* strains were cultured according to standard protocols [75]. Maintenance was done using Nematode Growth Medium (NGM) seeded with *E. coli* (OP50) at 20°C.

### Fertility and viability assays

Gravid adults of each genotype were bleached three consecutive times prior to the analysis. L4 stage hermaphrodites were individually picked into NGM plates seeded with a drop of OP50. Adults were transferred every 12 hr to new plates and the total number of laid eggs was scored. After 24 hours, the number of hatched progeny was scored.

### Lifespan assays

Lifespan assays were performed using regular NGM plates seeded with OP50 at 20°C, as previously described, but with some alterations [77]. Briefly, gravid adults were bleached three consecutive times prior to the assays. Twenty day 1 adult hermaphrodites were transferred to each NGM plate and denoted as “day 1”. Worms were scored as “censured” or “dead” every 2 to 3 days and transferred to fresh plates. Plates that became contaminated throughout the experiment were discarded. Worms were scored as “censured” when they crawled out of the plate, exhibited extruded intestine/vulva, or became “bags of worms”. Worms were scored as “dead” when their body no longer displayed movement in response to physical touch with a platinum wire and/or started to become transparent.

### Immunoblot analysis of whole-worm lysates

Gravid adults of each genotype were bleached for 3 consecutive rounds before collection. Day 1 adults were washed and collected in M9 buffer and snap frozen in liquid nitrogen. Worm pellets (50-100 µL of worms) were thawed on ice and 300 µL of Lysis buffer were added (Figure 1D, Lysis buffer composition: 50 mM Tris pH 7.5, 150 mM NaCl, 5 mM EDTA, 1% Triton X-100, 0.1% SDS, 1X Protease inhibitors, 25mM NEM, 25mM IAA, and 1mM PMSF; Figure 2G and Figure 6D, Lysis buffer composition: 50 mM Tris pH 7.5, 150 mM NaCl, 5 mM EDTA, 1% Triton X-100, 0.1% SDS, and 1X Protease inhibitors). Pellets were sonicated twice; each cycle was composed of 3X or 5X 5-second pulses (45% power) with 10-second intervals. Lysates were spun down (10000 rpm, 10 min) at 4°C and supernatant was collected. Protein concentration was determined using the BCA protein assay kit. Next, 40 µg of protein/sample were mixed with 5X laemmli buffer and heated at 95°C for 5 min prior to loading. Proteins were transferred to a nitrocellulose membrane for 90 to 180 min at 4°C. Membranes were washed with water for 5 min and blocked in 5% milk in 1X TBST for 1h at room temperature. Primary antibodies were incubated overnight in 5% milk in 1XTBST. Membranes were washed 3X with 1XTBST (10 min each) and subsequently incubated with HRP-conjugated secondary antibody (anti-mouse 1:10000, 1%milk in TBST, or anti-rabbit 1:10000, 1% milk in 1X TBST) for 1 hr at room temperature. For quantification, peak areas of the bands of proteins of interest (POI) were determined using gel tools in Fiji and normalized to the respective control bands. Fold change was determined by normalizing control samples to 1 and then determining the ratio of normalized POI in treated samples relative to the control samples.

## Antibodies

### RNA-seq sample preparation and analysis

Batches of 15 to 20 day 1 adults were transferred to a glass depression slide containing 10mM levamisole in M9. Worms were cut at the pharynx level with two syringe needles. Extruded gonads were separated from the worm body using scalpel blades, collected onto the forcep tip, and then transferred to eppendorfs containing 100 µL of trizol on ice. Up to 100 gonads were collected per 100 µL of trizol, then snap frozen in liquid nitrogen, and kept at -80°C for less than a week. A total of 250-270 gonads/sample were collected for the WT N2 strain and 320-340 gonads/sample for the F2 *ulp-2(tv380)* genotype. Total RNA was isolated using the Direct-zol RNA MiniPrep Plus kit following the manufacturer’s instructions (ZYMO RESEARCH). The RNA integrity number (RIN) was determined using TapeStation. Next, the Ultra II RNA Library prep kit was used to prepare the mRNA libraries and the HiSeq 2500 system was employed for sequencing. Adapters were trimmed with TrimGalore, read quality was inspected with FASTQC, and reads with less than 30bp were removed. After trimming, quality reads were mapped to the *C. elegans* reference genome with Tophat2, allowing a maximum of 3 mismatches/read. Uniquely mapped reads (more than 94% of the total reads) were counted using HTseq-count. Gene count normalization and differential expression analysis were performed with DESeq2 with a cut off value at p=0.05 (Benjamini-Hochberg correction). Gene Ontology analyses were performed using Panther (R. 20221013) with cut off at p>0.05 (FDR correction).

### Immunofluorescence

For Figure 2C, day 1 adults were transferred to a poly-lysine-coated slide in a drop of M9 and levamisole (10 mM). Two syringe needles were used to cut the worm at the pharynx level, followed by fixation with 1% PFA, then stained with DAPI (10 µg/µL) and mounted in Fluoromount-G. Z-stacks (0.3µm) of the germline were acquired using a Leica TCS SP5 II confocal microscope with a 63X 1.4 oil objective, along with the Leica LAS-AF software. The quantifications in Figures 2D and S1A-B were performed using the line tool and the multipoint tool in Fiji software. Confocal and DIC images in Figure 2E were acquired using a Zeiss confocal microscope with a 63X 1.4 oil objective.

### Yeast two hybrid

The Gal4-DB::ULP-2 bait construct was generated in vector pMB27, which was derived from vector pPC97 [79] by inserting an oligonucleotide linker that encodes a flexible linker (GGGG) upstream of the cloned ORF, and enables cloning using the AscI and NotI restriction sites. The oligonucleotide linker was ligated into pPC97 digested with SmaI and SacI. The ULP-2 bait fragment used consists of the first 5 coding exons (bp 4-771). The corresponding *ulp-2* sequence was amplified from cDNA by PCR and cloned into pMB27 digested with AscI and NotI.

The Gal4-DB::ULP-2 fusion plasmid was transformed into *S. cerevisiae* strain Y8930 [80], and then mated to a pPC86-Gal4-AD prey library of mixed-stage *C. elegans* cDNAs transformed into *S. cerevisiae* strain Y8800 [80] (a gift from Xiaofeng Xin and Charlie Boone). Yeast expressing putative interacting protein pairs were selected by plating mated yeast on synthetic complete (SC) medium plates lacking leucine, tryptophan, and histidine. At least 2x10^6^ independent colonies were screened. *De novo* autoactivating yeast colonies were eliminated by a plasmid-shuffling-based counter selection [81]. The identities of candidate interacting proteins, which included SET-26, were then determined by PCR amplification and sequencing of the cDNA inserts.

To confirm the interaction between ULP-2 and SET-26, an N-terminal fragment (residues 1–575) and a C-terminal fragment of SET-26 (residues 1188–1645) were cloned into Gal4-AD vector pMB29 [82], transformed into *S. cerevisiae* strain Y8800, mated with strain Y8930 expressing Gal4-DB::ULP-2, and then assayed for protein interaction on selective SC agar plates lacking leucine, tryptophan, and histidine or adenine. Plates lacking histidine were supplemented with 2 mM 3-Amino-1,2,4-triazole (3-AT). In addition, auto-activation was assayed on an SC plate lacking leucine and histidine, supplemented with 1µg/ml cycloheximide.

### RNA interference (RNAi)

RNAi was performed by dsRNA injection for the experiments shown in Figure S3B-E. dsRNA from exons 1-4 and exon 15 of *ulp-2* and exons 5-7 of Y73Ba5.1/SET-9/SET-26 (“Set3”) were synthetized in vitro with the T7 RiboMAX express kit or the HiScribe T7 high-yield RNA synthesis kit following the manufacturer’s instructions. All dsRNA were injected at a concentration of 50 ng/µL. For Figures 6A and S5B-C, RNAi was performed by feeding as previously described [36]. In brief, HT115 bacteria were transformed with two *ulp-2(RNAi)* clones (exons 1-4 and exon 15) and L4440 control and grown overnight at 37°C. A minimum of 10 colonies of each RNAi clone were grown in 2 mL LB+ampicillin overnight and diluted 1:100 with fresh LB+ampicillin medium for an additional 4-6h until OD600=1. The two *ulp-2(RNAi)* bacterial pellets were concentrated 10 times with M9 and mixed in a 1:1 ratio. Aliquots of 800 µL were seeded in 20 mL NGM plates containing 1 mM IPTG and 25 µg/ml carbenicillin and dried overnight. IPTG (200 µL, 100 mM stock) was spread above the bacterial lawn and allowed to dry before embryos were dropped.

### In-vitro sumoylation and desumoylation

The SET domain (AA 945-1107) or the PHD-SET (AA 794-1107) domains of SET-26 were cloned into pGEX-4T1 and transformed into *E. coli* BL21(DE3)pLysS. Colonies were grown for 2.5 hr at 37°C and then induced with 0.1mM isopropyl β-D-1-thiogalactopyranoside (IPTG) for 4 hr at 37°C. Proteins were purified with Lysis buffer (50mM Tris pH 7.5, 150 mM NaCl, 0.05% NP-40, EDTA-free proteinase inhibitor, 1mM PMSF, and 250 µg/ml Lysozyme). Proteins were affinity purified on glutathione resin (GeneScript #L00206) and eluted with 100 mM Tris pH 8.0 and 10 mM reduced glutathione (Sigma, #G4251). After protein determination, 2.5 µg protein were used for a sumoylation reaction performed according to the manufacturer’s protocol (LAE Biotech). In-vitro sumoylation reactions were incubated at 37°C for the indicated time points.

For the in-vitro desumoylation reactions, the ULP-2 catalytic domain (AA 501-894) was cloned into pET28a+ and transformed into *E. coli* BL21(DE3). Colonies were grown for 2 hr at 37°C (OD600 ∼0.8) and then induced with 1 mM isopropyl β-D-1-thiogalactopyranoside (IPTG) for 4 h at 30°C. Proteins were purified with Lysis buffer (20% Sucrose, 20 mM Tris pH8, 1mM β-mercaptoethanol, 350 mM NaCl, 20 mM Imidazole, 10 µg/ml DNase, 1 mM PMSF, 0.1% Igepal CA630, 20 µg/ml Lysozyme, and EDTA-free proteinase inhibitor). Proteins were affinity purified from lysate by Ni Sepharose 6 Fast Flow resin (GE Healthcare #17-5318-01) and eluted with 20 mM Tris pH8, 1mM β-mercaptoethanol, 350 mM NaCl, and 400 mM Imidazole. Then, they were concentrated using Amicon centrifugal filters 10,000 NMWL. Deconjugation reactions were performed in 25mM Tris pH8, 150mM NaCl, 0.1% Tween, and 2mM DTT according to [83], and ∼5 ng ULP-2. The PHD-SET domain was sumolyated for 2 hr before deconjugation at 37°C for the indicated time points.

### In-vitro methylation

The methylation assay reactions contained a pre-sumoylated or non-sumoylated GST-tagged PHD-SET domain of SET-26, 2 mCi of 3H-labeled S-adenosyl-methionine (SAM) (Perkin-Elmer, AdoMet), and PKMT buffer (20 mM Tris–HCl pH 8, 10% glycerol, 20 mM KCl, and 5 mM MgCl_2_). The reactions were incubated overnight at 30°C and were then resolved by SDS-PAGE for Coomassie staining (Expedeon InstantBlue) or autoradiography.

### Immunoprecipitation

Samples were mainly collected as described above in “immunoblot analysis” with the following exception: the worm pellets were incubated with 1% of PFA for 15 min at room temperature, washed 2Xs with 50 mM Tris pH=8 (5 min), and subsequently snap frozen in liquid nitrogen. Samples were thawed on ice and 300 µL of Lysis buffer (50 mM Tris pH 7.5, 150 mM NaCl, 5 mM EDTA, 1% Triton X-100, 0.1% SDS, 1.3X Protease inhibitors, and 25mM NEM) were added. Samples were sonicated on ice in two cycles as indicated above and subsequently spun down at 7000 rpm, 5 min at 4°C. Next, 35 µL of supernatant were reserved for “input” and the remnant was pre-cleared with 15-25 µL of pre-washed Protein G Sepharose beads (GE healthcare, 17-0618-01) (3X, 5 min, at 4°C) for 30 min at 4°C with rotation. Beads were spun down for 20 sec at 12000 rpm and the supernatant was transferred to an eppendorf tube containing 8-10 µL of anti-GFP antibody and incubated on ice for 1 h. Next, 30-50 µL of pre-washed beads were added and samples were incubated at 4°C for 1 h with rotation. Antibody-bead conjugates were washed once with Lysis buffer and 3X with 50 mM Tris pH=8 (10 min) (in Figure 6A, the antibody-bead conjugates were washed 2X with Lysis buffer). Finally, 5X laemmli buffer was added and heated at 95°C for 2 min. The primary antibodies used were anti-GFP, anti-H3K4me3, and anti-sumo (Table 2).

**Table 1.**
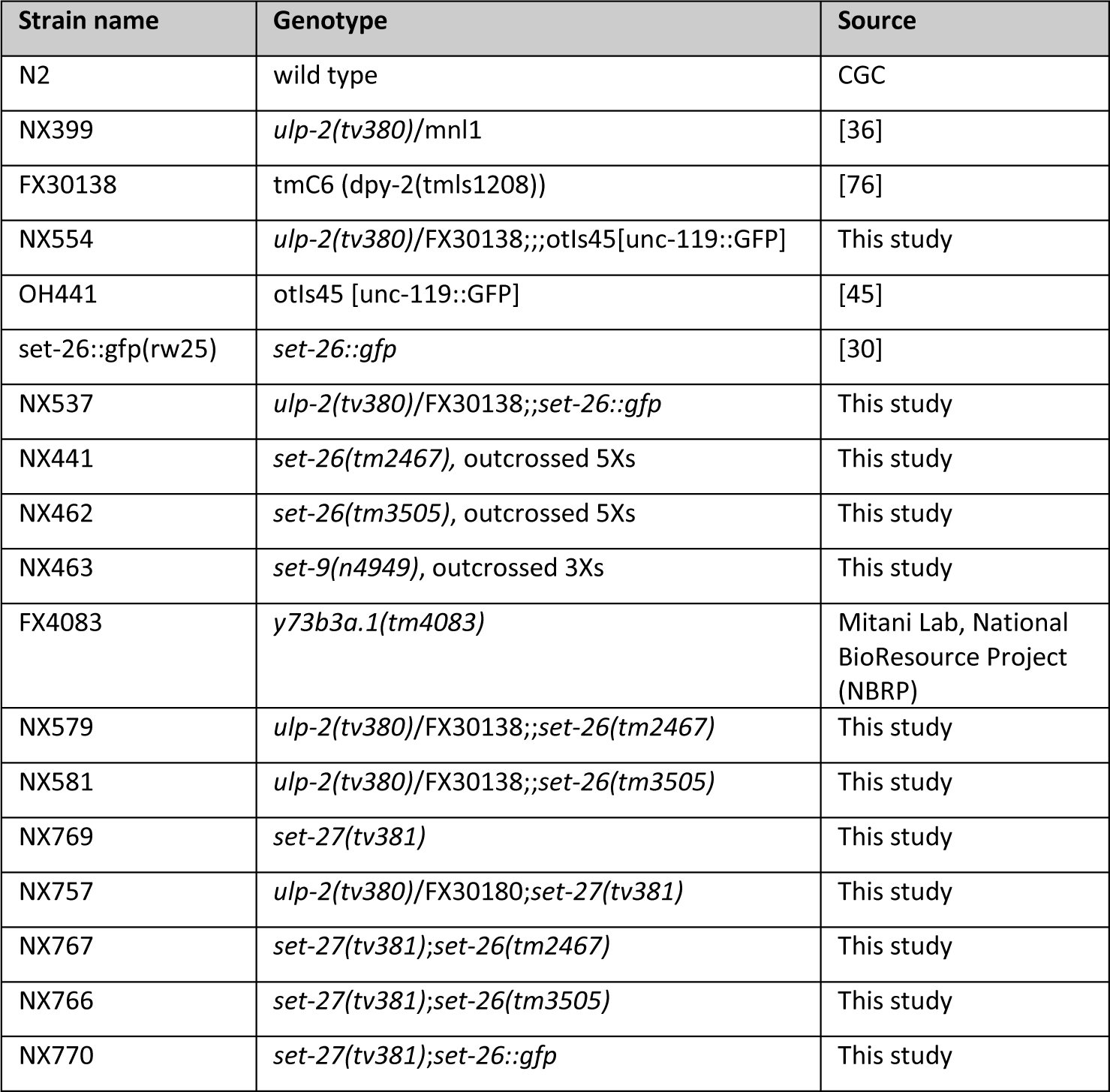
Strains used in this study.

**Table 2.** Antibodies and dilutions used in this study.

| Antibody (dilution) | Catalog #, Company |
| --- | --- |
| anti-sumo (1:160) | SUMO 6F2, DSHB [78] |
| anti-α tubulin DM1A (1:1000) | 3873, CST |
| anti-H3 (1:10,000) | ab1791, Abcam |
| anti-H3 (1:1000) | ab24834, Abcam |
| anti-H3K4me3 (1:2000-5000) | ab8580, Abcam |
| anti-H3K27me3 (1:1000) | ab192985, Abcam |
| anti-GFP (1:5000-7000) | 11814460001, Roche |
| anti-GST (B-14) (1:250) | SC-138, Santa Cruz Biotechnology |

### Label-free mass spectrometry

N2 and SET-26::GFP day 1 adults were bleached and embryos were dropped on 20 cm NGM plates containing IPTG and carbenicillin for RNAi by feeding (L4440 and *ulp-2* clones) as described above. Following RNAi feeding, adult worms were collected and immunoprecipitation was carried out as described above. Immunoprecipitants were loaded on a 10% acrylamide gel and separated 3cm within the gel. Gels were stained with Coomassie blue overnight and destained with destaining solution for 3-4 hours. The proteins in the gel were reduced with 3mM DTT (at 60°C for 30 min), modified with 10mM iodoacetamide in 100 mM ammonium bicarbonate (in the dark, at room temperature for 30 min) and then digested in 10% acetonitrile and 10 mM ammonium bicarbonate with trypsin at a 1:10 enzyme-to-substrate ratio, overnight at 37°C. The resulting peptides were desalted using C18 tips (homemade stage tips) and were subjected to LC-MS-MS analysis. The peptides were resolved by reverse-phase chromatography on 0.075 X 180-mm fused silica capillaries packed with ReproSil reversed phase material. The peptides were eluted with a linear 60-minute gradient of 5 to 28%, a 15-minute gradient of 28 to 95%, and a 25-minute gradient using 95% acetonitrile with 0.1% formic acid in water at flow rates of 0.15 μl/min. Mass spectrometry was performed using a Q Exactive HF mass spectrometer in a positive mode (m/z 300–1800, resolution 120,000 for MS1, and 15,000 for MS2) using repetitively a full MS scan, followed by collision induced dissociation (HCD, at 27 normalized collision energy) of the 18 most dominant ions (>1 charges) selected from the first MS scan. The AGC settings were 3x106 for the full MS and 1x105 for the MS/MS scans. The intensity threshold for triggering MS/MS analysis was 1x104. A dynamic exclusion list was enabled with an exclusion duration of 20s. The mass spectrometry data were analyzed using MaxQuant software 1.5.2.8 for peak picking and identification using the Andromeda search engine, searching against the *Caenorhabditis elegans* proteome from the Uniprot database with a mass tolerance of 6 ppm for the precursor masses and 20 ppm for the fragment ions. Oxidation on methionine and protein N-terminus acetylation were accepted as variable modifications, and carbamidomethyl on cysteine was accepted as static modifications. The minimal peptide length was set to six amino acids and a maximum of two miscleavages was allowed. The data were quantified by label-free analysis using the same software. Peptide- and protein-level false discovery rates (FDRs) were filtered to 1% using the target-decoy strategy. The protein table was filtered to eliminate the identifications from the reverse database, as well as common contaminants and single peptide identifications. The imputation of missing values was set at 20 and peptide intensities were normalized to the peptide’s intensities of the SET-26 protein in each sample. Statistical analysis of the identification and quantization results was done using Perseus software (1.6.10.43). Gene ontology analysis was performed in Panther.

### CRISPR/Cas9 genome editing

Generation of the genome-edited *set-27(tv381)* strain was performed using CRISPR/Cas9 technology adapted from [84]. The injection mix was prepared by mixing 1 µl of Cas9 (IDT), 5 µl of tracrRNA, 1 µl dpy-10 crRNA, 2 µl of *set-27(tv381)* crRNA-1, and 2 µl of crRNA-2 and incubated at 37°C for 20 min. Next, 2 µl of *dpy-10* ssODN, 4 µl of *set-27* knockout repair primer, and 3 µl of ddH2O were added (Table S5). The mix was spun down for 2 min at 12000 rpm. Then, 17 µl of the mix were transferred to a new PCR tube and injected into the gonads of N2 day 1 adult worms. P0 animals were transferred individually to new plates after ∼12 h. Dpy or roller phenotypes were screened to identify P0 worms with successful Cas9 delivery into their germline. F1 progeny (dpy, roller, or non-dpy) of successfully injected P0 was individually isolated to new plates and allowed to lay for ∼2 days before genotyping.

### Quantification and statistical analyses

All statistical analysis were performed with GraphPad Prism (version 10.0.2) except for RNA-seq and Mass Spectrometry data as described above.

## Acknowledgments

This research was supported by the Israel Science Foundation ISF grants 2122/19 (personal research grant and a grant for international cooperation with MB) and 1878/15 to LB. Several strains were provided by the CGC, which is funded by the NIH Office of Research Infrastructure Programs (P40 OD010440) and by the National Bioresource Project, Tokyo, Japan NBRP, which is funded by the Japanese government. The SET-26::GFP strain was sent to us from the Siu Sylvia Lee laboratory at Cornell Univ., Ithaca, NY, USA. RNA-seq was performed at the Technion Genomics Center and Proteomics was performed at The Technion Proteomics Center, Haifa, Israel.

## Author contributions

**Cátia A. Carvalho:** Conceptualization, Formal analysis, Investigation, Methodology, Project administration, Visualization, Writing-original draft, Writing-review&editing. **Ulrike Bening Abu-Shach:** Investigation. **Asha Raju:** Investigation. **Zlata Vershinin:** Investigation. **Dan Levy:** Methodology, Writing-original draft, Writing-review&editing. **Mike Boxem:** Investigation, Methodology, Funding acquisition, Writing-original draft, Writing-review&editing. **Limor Broday:** Conceptualization, Formal analysis, Funding acquisition, Investigation, Methodology, Project administration, Resources, Supervision, Validation, Visualization, Writing-original draft, Writing-review&editing.

## Disclosure and competing interests statement

The authors declare no competing interests.

## Supplementary Figures and Tables

**Figure S1.**
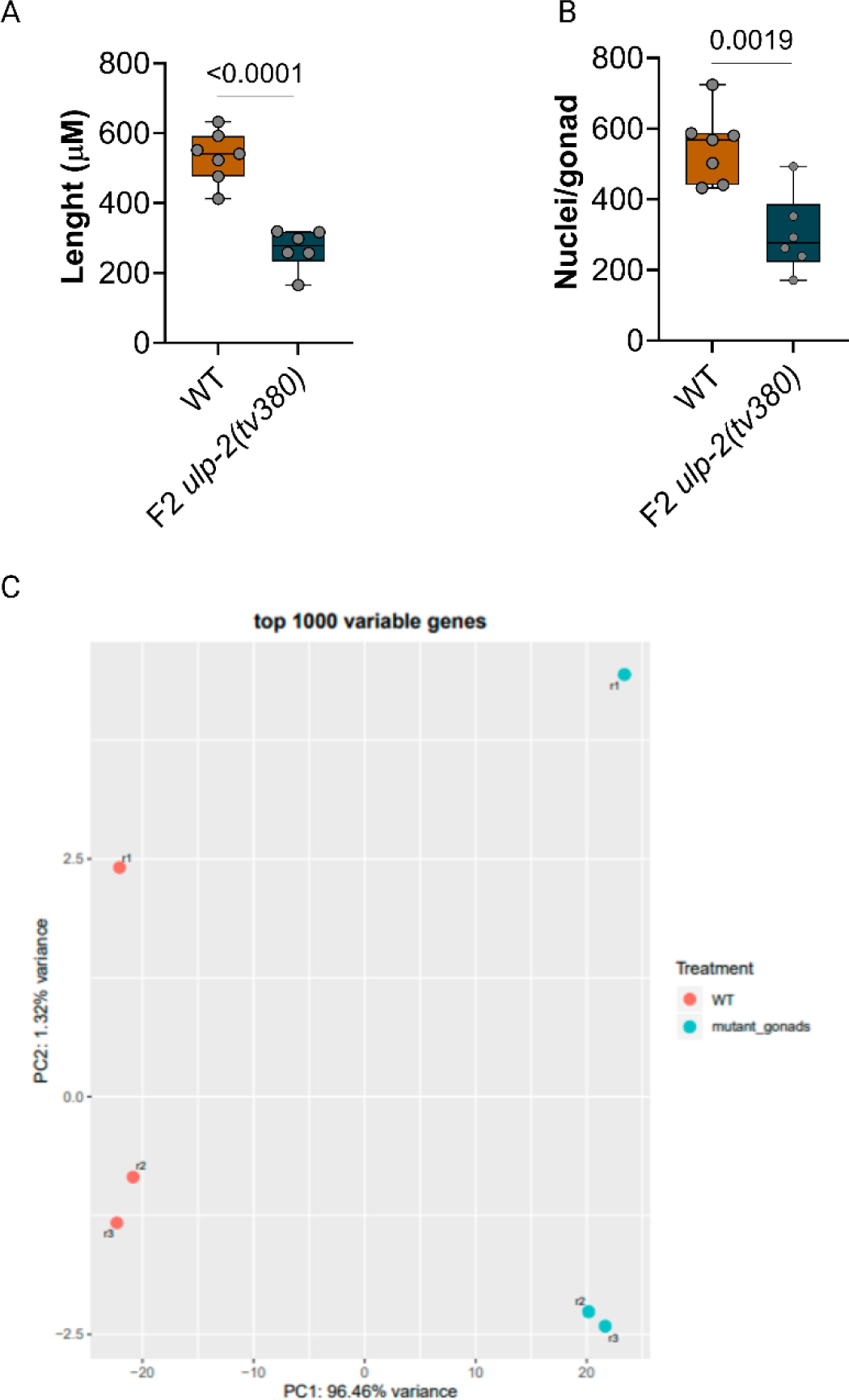
ULP-2 loss of function reduces the length and cell content in the germline and disrupts gene expression. **A.** Quantification of germline length of WT (7 germlines) and F2 *ulp-2(tv380)* (6 germlines); Two-tailed Welch’s t-test was used, ns=p>0.05; **B.** Quantification of the total number of nuclei in WT (3834 nuclei) and F2 *ulp-2(tv380)* (1809 nuclei); Two-tailed Welch’s t-test was used, ns=p>0.05; **C.** PCA analysis of the RNA-seq dataset of WT (3 biological samples, 779 isolated germlines) and F2 *ulp-2(tv380)* (3 biological samples, 990 isolated germlines); pink dots represent the biological replicates of WT isolated germlines and blue dots represent the biological replicates of F2 *ulp-2(tv380)* isolated germlines.

**Figure S2.**
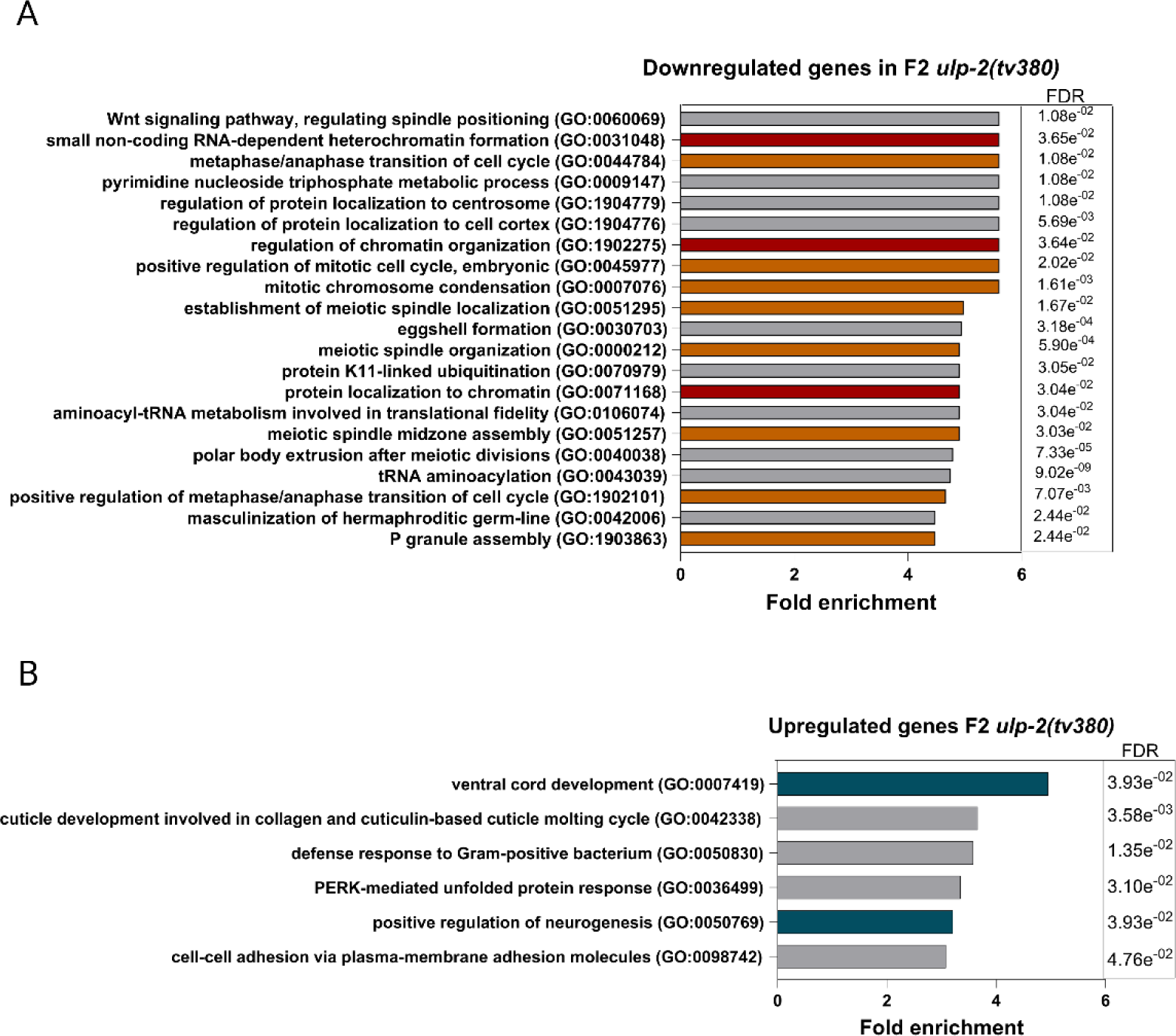
ULP-2 loss of function induces downregulation of genes involved in germline development and upregulation of genes involved in somatic functions in the germline. **A.** Gene ontology analysis of the biological processes containing the downregulated genes in germlines of F2 *ulp-2(tv380)* when compared to WT; Biological processes were cut off at Fold enrichment<4; ns=FDR>0.05. **B.** Gene ontology analysis of the biological processes containing the significantly upregulated genes in F2 *ulp-2(tv380)* when compared to WT; Biological processes were cut off at Fold enrichment<2.5; ns=FDR>0.05. GO terms mentioned in the manuscript text appear in color.

**Figure S3.**
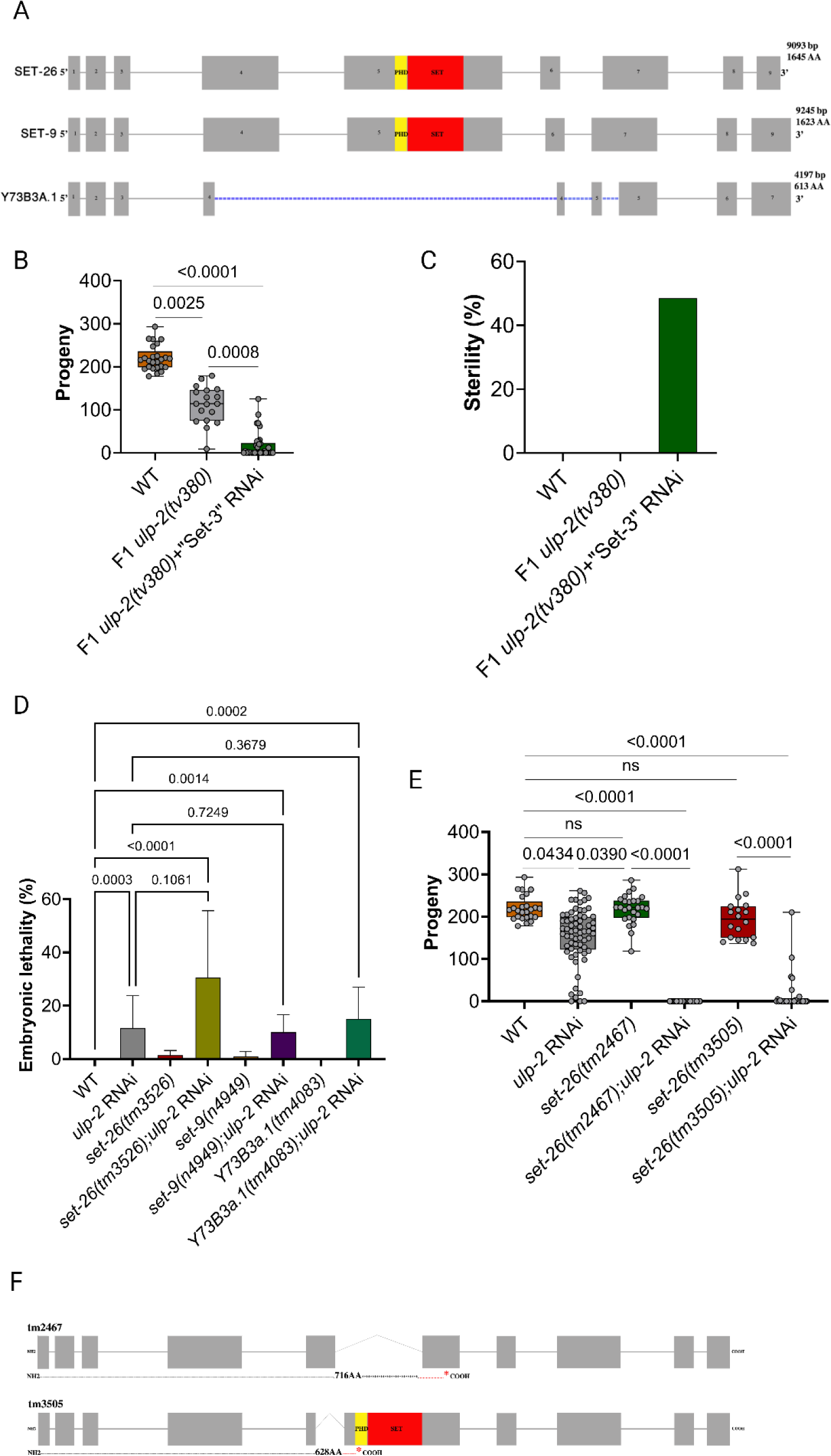
SET-26 interacts genetically with ULP-2 in *C. elegans*. **A.** Schematic representation of the exons composing the three family members of the predicted methyltransferase family identified in the Y2H screen; SET and PhD domains encoding exons are highlighted in red and yellow, respectively; **B.** Quantification of brood size of WT (25 worms), first generation of *ulp-2(tv380)* mutant animals (18 worms) and concomitant downregulation of the three family members (*ulp-2(tv380)*+”Set3” RNAi) (34 worms); Shapiro-Wilk and One-Way Anova on ranks (Kruskal-Wallis) followed by Dunn’s post-hoc test was used, ns=p>0.05. **C.** Percentage of quantified mothers in B. that exhibited sterility. **D.** Quantification of embryonic lethality of WT (8 worms), *ulp-2*(RNAi) (32 worms), *set-26(tm3526)* (4 worms), *set-26(tm3526)*;*ulp-2(RNAi)* (6 worms), *set-9(n4949)* (3 worms), *set-9(n4949)*;*ulp-2(RNAi)* (9 worms), Y73B3a.1(tm4083) (3 worms) and Y73B3a.1(tm4083);*ulp-2(RNAi)* (10 worms); Shapiro-Wilk and One-Way Anova on ranks (Kruskal-Wallis) followed by Dunn’s post-hoc test was used, ns=p>0.05. **E.** Quantification of amount of progeny of WT (25 worms), *ulp-2(RNAi)* (65 worms), *set-26(tm2467)* (26 worms), *set-26(tm2467)*;*ulp-2(RNAi)* (80 worms), *set-26(tm3505)* (18 worms) and *set-26(tm3505);ulp-2(RNAi)* (30 worms); Shapiro-Wilk and One-Way Anova on ranks (Kruskal-Wallis) followed by Dunn’s post-hoc test was used, ns=p>0.05. **F.** Schematic representation of WT and *set-26* deletion alleles (NBRP, Japan) *set-26(tm2467)* and *set-26(tm3505),* early stop codon is labeled in red.

**Figure S4.**
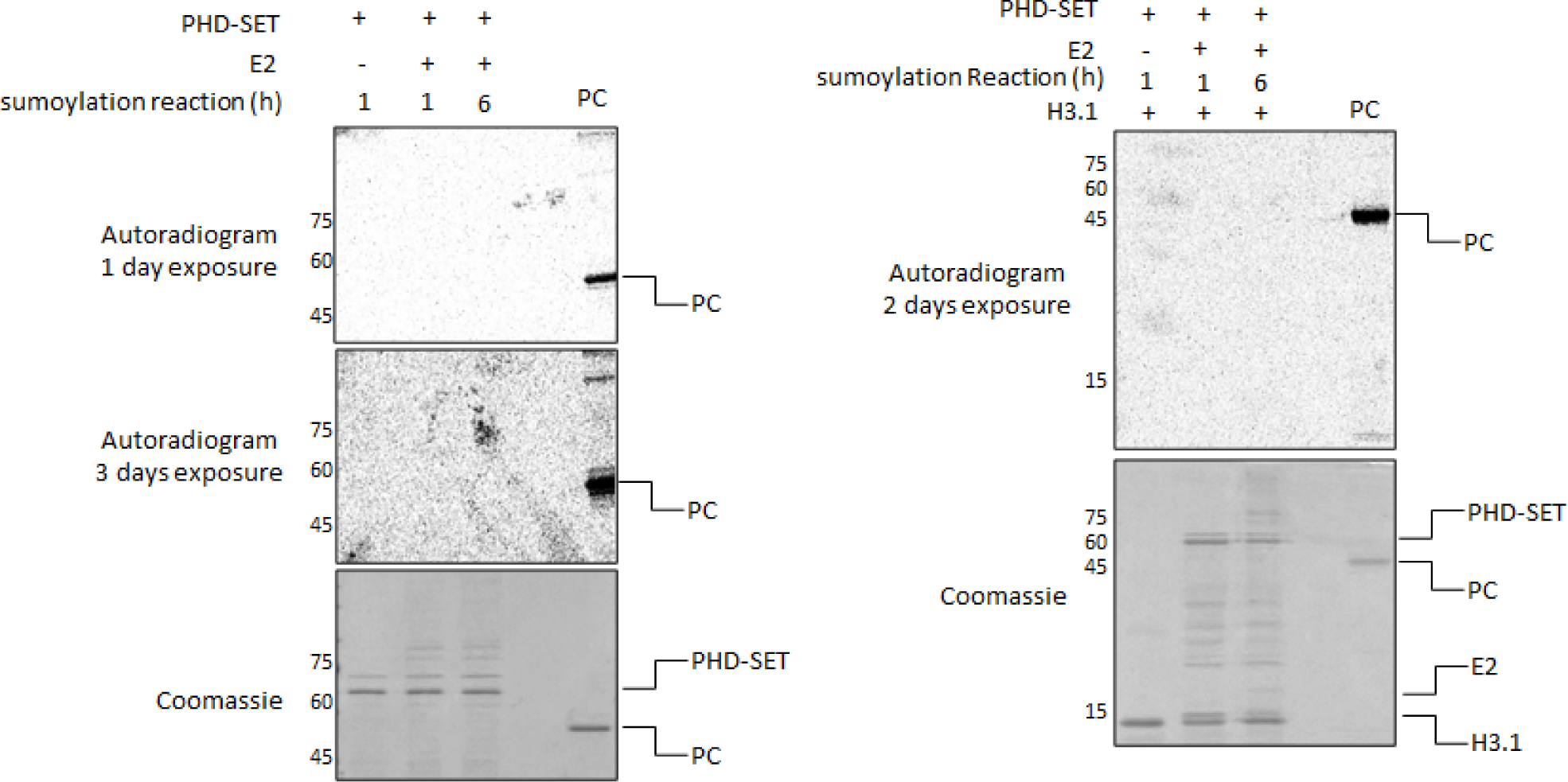
The PHD-SET domain of SET-26 is enzymatically inactive. **(A+B)** Recombinant GST-PHD-SET of SET-26 was sumoylated for the indicated time points with or without the addition of E2 (UBC9). Samples were then subjected to an in-vitro methylation reaction in the presence of 3H-labeled SAM without (A) or with recombinant Histone H3.1 (H3.1) as a substrate (B). Samples were subjected to SDS–polyacrylamide gel electrophoresis (PAGE) followed by exposure to autoradiogram as indicated. Human SETD6 served as positive control (PC). Coomassie stain of the recombinant proteins used in the reactions is shown on the bottom.

**Figure S5.**
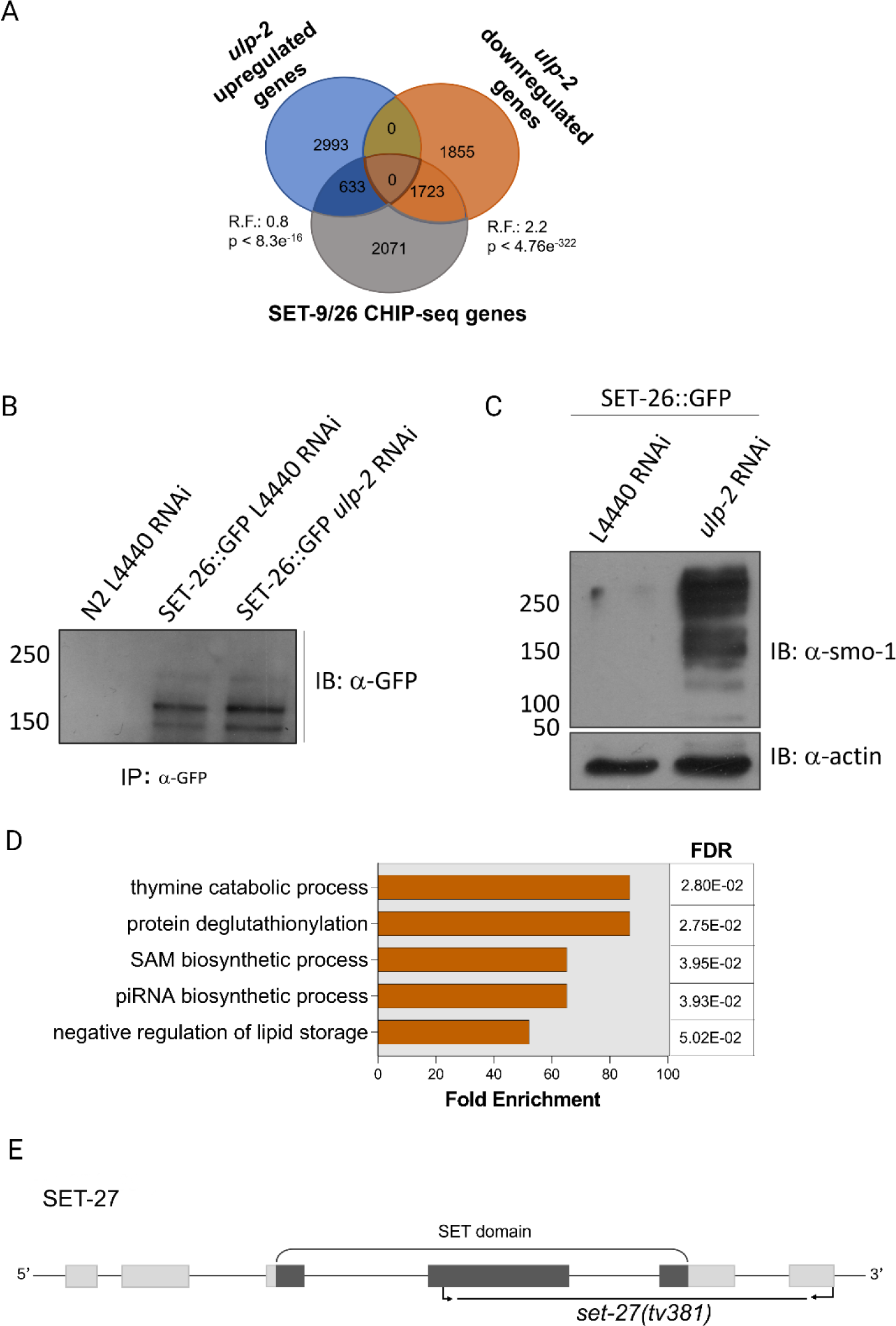
Downregulation of ULP-2 weakens SET-26 complex formation. **A.** Venn diagram representing intersection between the differential expressed genes in F2 *ulp-2(tv380)* germlines and the SET-9/26 binding genes (CHIP-seq); Fischer’s exact test was used, ns=p>0.05, R.F.=Representation factor. **B.** Representative immunoblot of the IP of SET-26::GFP samples sent to Mass Spectrometry; N2 L4440 RNAi was used as a control for GFP immunoprecipitation, SET-26::GFP L4440 RNAi corresponds to SET-26::GFP in a WT background and SET-26::GFP *ulp-2(RNAi)* corresponds to SET-26::GFP IP in the excessive SUMOylation background; 3 biological replicates. **C.** Representative immunoblot showing excessive SUMOylation in the knockdown of ULP-2 *ulp-2(RNAi)* when compared to the control (L4440 RNAi). **D.** Gene Ontology analysis of the WT interacting partners of SET-26::GFP. **E.** Exon intron structure of SET-27 and *set-27(tv381)* deletion allele. The light grey rectangles and dark rectangles are exons, the dark part of the rectangles is the SET domain, the arrows is the place where the crRNAs bind and the deletion is highlighted.

**Figure S6.**
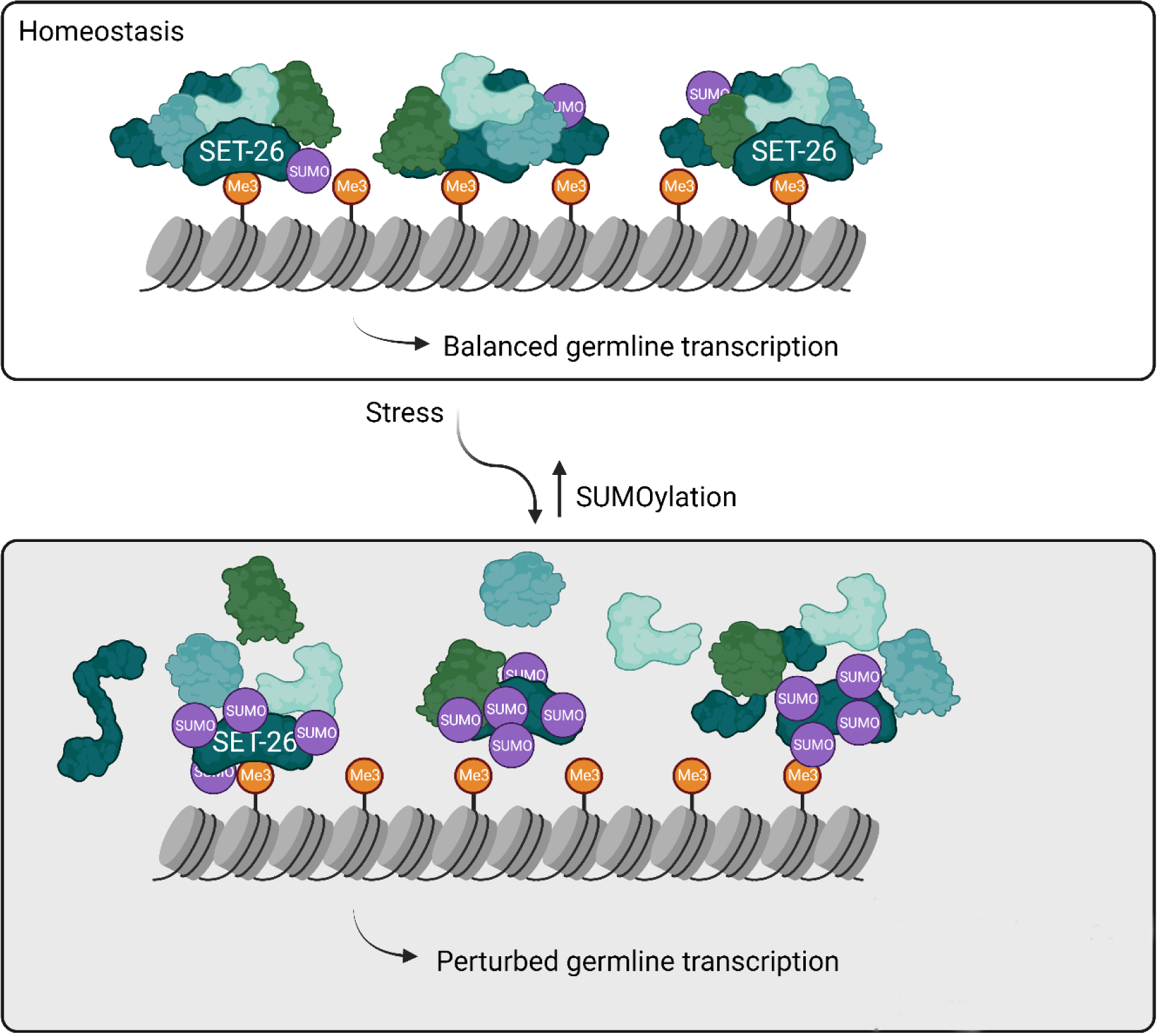
Model for SET-26 dependence on sumoylation. In homeostasis, SET-26 and its interactors bind H3K4me3 marks to regulate gene expression. Unbalanced sumoylation of SET-26 disrupts its reader function by decreasing its binding to H3K4me3 marks. Consequently, germline transcription is perturbed.

**Table S1.** Lifespan analysis at 20°C.

| Genotype | No. of deaths | Median Lifespan (days) | P-value vs. N2 | P-value vs. F1 *ulp-2(tv380)* | P-value vs. F2 *ulp-2(tv380)* |
| --- | --- | --- | --- | --- | --- |
| N2 | 252 | 16 | 1 | $< 10^{-3}$ | $< 10^{-3}$ |
| F1 *ulp-2(tv380)* | 220 | 19 | $< 10^{-3}$ | 1 | $< 10^{-3}$ |
| F2 *ulp-2(tv380)* | 213 | 24 | $< 10^{-3}$ | $< 10^{-3}$ | 1 |
| *set-26(tm2467)* | 254 | 23 | $< 10^{-3}$ | $< 10^{-3}$ | 0.0188 |
| *ulp-2(tv380);set-26(tm2467)* | 157 | 18 | $< 10^{-3}$ | 0.4577 | $< 10^{-3}$ |
| *set-26(tm3505)* | 245 | 25 | $< 10^{-3}$ | $< 10^{-3}$ | 0.1672 |
| *ulp-2(tv380);set-26(tm3505)* | 205 | 18 | 0.1154 | 0.0222 | $< 10^{-3}$ |

**Table S5.**
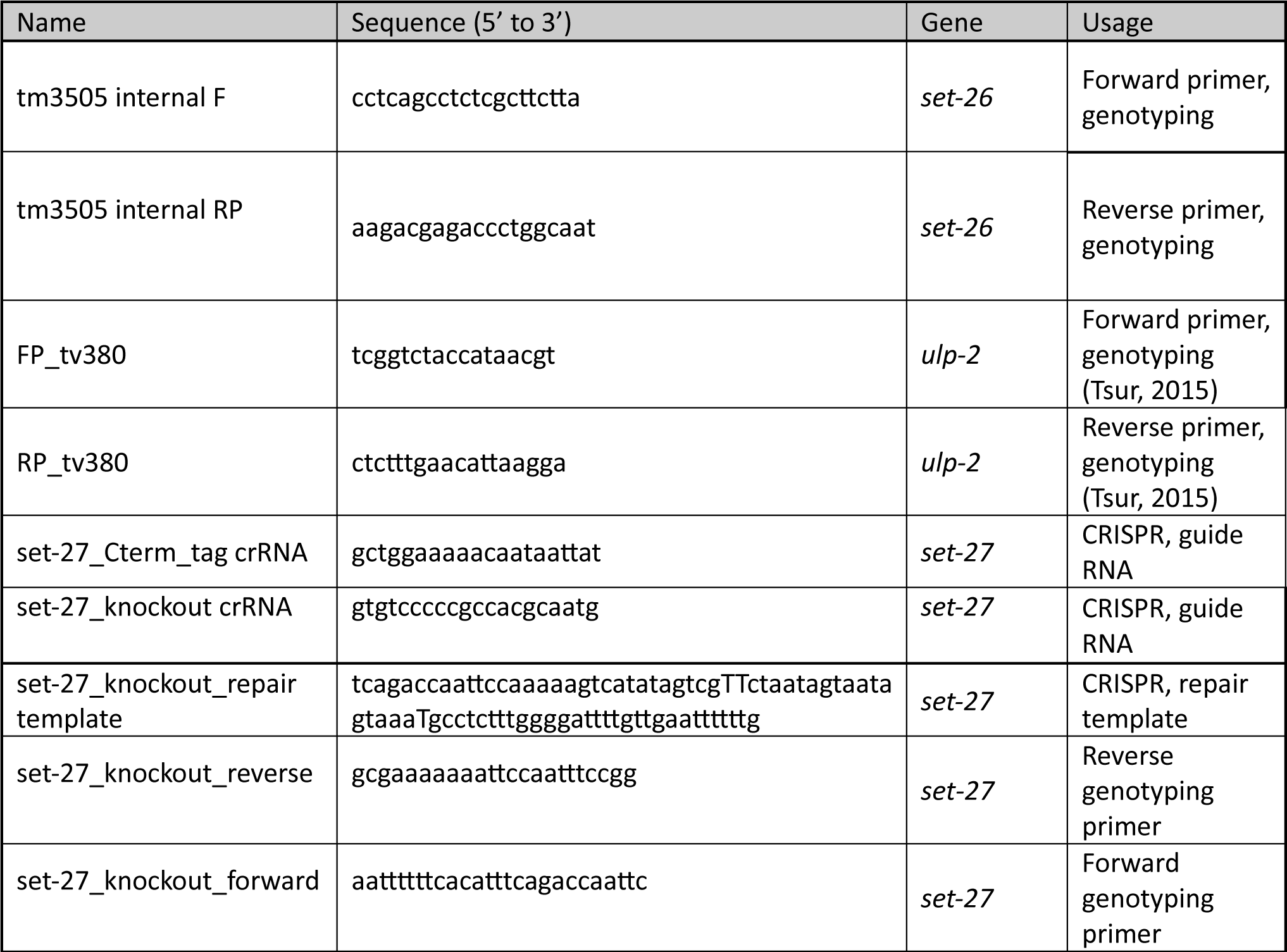
List of PCR primers, gRNA and ssOligo donor sequences.

## Notes

### Competing Interest Statement

The authors have declared no competing interest.

## References

1. Matunis MJ, Coutavas E, Blobel G: A novel ubiquitin-like modification modulates the partitioning of the Ran-GTPase-activating protein RanGAP1 between the cytosol and the nuclear pore complex. J Cell Biol 1996, 135:1457–1470.

2. Mahajan R, Delphin C, Guan T, Gerace L, Melchior F: A small ubiquitin-related polypeptide involved in targeting RanGAP1 to nuclear pore complex protein RanBP2. Cell 1997, 88:97–107.

3. Flotho A, Melchior F: Sumoylation: A Regulatory Protein Modification in Health and Disease. Annual Review of Biochemistry 2013, 82:357–385.

4. Mukhopadhyay D, Dasso M: Modification in reverse: the SUMO proteases. Trends in Biochemical Sciences 2007, 32:286–295.

5. Hickey CM, Wilson NR, Hochstrasser M: Function and regulation of SUMO proteases. Nature Reviews Molecular Cell Biology 2012, 13:755–766.

6. Chymkowitch P, Nguéa PA, Enserink JM: SUMO-regulated transcription: challenging the dogma. Bioessays 2015, 37:1095–1105.

7. Hendriks IA, Treffers LW, Verlaan-de Vries M, Olsen JV, Vertegaal ACO: SUMO-2 Orchestrates Chromatin Modifiers in Response to DNA Damage. Cell Rep 2015, 10:1778–1791.

8. Hendriks IA, Vertegaal AC: A comprehensive compilation of SUMO proteomics. Nat Rev Mol Cell Biol 2016, 17:581–595.

9. Vertegaal ACO: Signalling mechanisms and cellular functions of SUMO. Nat Rev Mol Cell Biol 2022, 23:715-731.

10. Strahl BD, Allis CD: The language of covalent histone modifications. Nature 2000, 403:41–45.

11. Jenuwein T, Allis CD: Translating the histone code. Science 2001, 293:1074–1080.

12. Kang X, Qi Y, Zuo Y, Wang Q, Zou Y, Schwartz RJ, Cheng J, Yeh ETH: SUMO-Specific Protease 2 Is Essential for Suppression of Polycomb Group Protein-Mediated Gene Silencing during Embryonic Development. Molecular Cell 2010, 38:191–201.

13. Zhang H, Smolen GA, Palmer R, Christoforou A, van den Heuvel S, Haber DA: SUMO modification is required for in vivo Hox gene regulation by the Caenorhabditis elegans Polycomb group protein SOP-2. Nat Genet 2004, 36:507–511.

14. Kagey MH, Melhuish TA, Wotton D: The polycomb protein Pc2 is a SUMO E3. Cell 2003, 113:127–137.

15. Cossec JC, Theurillat I, Chica C, Búa Aguín S, Gaume X, Andrieux A, Iturbide A, Jouvion G, Li H, Bossis G, Seeler JS, Torres-Padilla ME, Dejean A: SUMO Safeguards Somatic and Pluripotent Cell Identities by Enforcing Distinct Chromatin States. Cell Stem Cell 2018, 23:742–757.e748.

16. Ninova M, Godneeva B, Chen YA, Luo Y, Prakash SJ, Jankovics F, Erdélyi M, Aravin AA, Fejes Tóth K: The SUMO Ligase Su(var)2-10 Controls Hetero- and Euchromatic Gene Expression via Establishing H3K9 Trimethylation and Negative Feedback Regulation. Mol Cell 2020, 77:571–585.e574.

17. Ruthenburg AJ, Li H, Patel DJ, Allis CD: Multivalent engagement of chromatin modifications by linked binding modules. Nat Rev Mol Cell Biol 2007, 8:983–994.

18. Bannister AJ, Zegerman P, Partridge JF, Miska EA, Thomas JO, Allshire RC, Kouzarides T: Selective recognition of methylated lysine 9 on histone H3 by the HP1 chromo domain. Nature 2001, 410:120–124.

19. Lachner M, O’Carroll D, Rea S, Mechtler K, Jenuwein T: Methylation of histone H3 lysine 9 creates a binding site for HP1 proteins. Nature 2001, 410:116–120.

20. Baytek G, Blume A, Demirel FG, Bulut S, Popp O, Mertins P, Tursun B: SUMOylation of the chromodomain factor MRG-1 in C. elegans affects chromatin-regulatory dynamics. Biotechniques 2022, 73:5–17.

21. Sanchez R, Zhou MM: The PHD finger: a versatile epigenome reader. Trends Biochem Sci 2011, 36:364–372.

22. Li H, Ilin S, Wang W, Duncan EM, Wysocka J, Allis CD, Patel DJ: Molecular basis for site-specific read-out of histone H3K4me3 by the BPTF PHD finger of NURF. Nature 2006, 442:91–95.

23. Peña PV, Davrazou F, Shi X, Walter KL, Verkhusha VV, Gozani O, Zhao R, Kutateladze TG: Molecular mechanism of histone H3K4me3 recognition by plant homeodomain of ING2. Nature 2006, 442:100–103.

24. Ivanov AV, Peng H, Yurchenko V, Yap KL, Negorev DG, Schultz DC, Psulkowski E, Fredericks WJ, White DE, Maul GG, Sadofsky MJ, Zhou MM, Rauscher FJ, 3rd: PHD domain-mediated E3 ligase activity directs intramolecular sumoylation of an adjacent bromodomain required for gene silencing. Mol Cell 2007, 28:823–837.

25. Madan V, Madan B, Brykczynska U, Zilbermann F, Hogeveen K, Döhner K, Döhner H, Weber O, Blum C, Rodewald HR, Sassone-Corsi P, Peters AH, Fehling HJ: Impaired function of primitive hematopoietic cells in mice lacking the Mixed-Lineage-Leukemia homolog MLL5. Blood 2009, 113:1444–1454.

26. Zhou P, Wang Z, Yuan X, Zhou C, Liu L, Wan X, Zhang F, Ding X, Wang C, Xiong S, Wang Z, Yuan J, Li Q, Zhang Y: Mixed lineage leukemia 5 (MLL5) protein regulates cell cycle progression and E2F1-responsive gene expression via association with host cell factor-1 (HCF-1). J Biol Chem 2013, 288:17532–17543.

27. Lemak A, Yee A, Wu H, Yap D, Zeng H, Dombrovski L, Houliston S, Aparicio S, Arrowsmith CH: Solution NMR structure and histone binding of the PHD domain of human MLL5. PLoS One 2013, 8:e77020.

28. Ali M, Rincón-Arano H, Zhao W, Rothbart SB, Tong Q, Parkhurst SM, Strahl BD, Deng LW, Groudine M, Kutateladze TG: Molecular basis for chromatin binding and regulation of MLL5. Proc Natl Acad Sci U S A 2013, 110:11296–11301.

29. Tran K, Green EM: SET domains and stress: uncovering new functions for yeast Set4. Curr Genet 2019, 65:643–648.

30. Wang W, Chaturbedi A, Wang M, An S, Santhi Velayudhan S, Lee SS: SET-9 and SET-26 are H3K4me3 readers and play critical roles in germline development and longevity. Elife 2018, 7.

31. Pijnappel WW, Schaft D, Roguev A, Shevchenko A, Tekotte H, Wilm M, Rigaut G, Séraphin B, Aasland R, Stewart AF: The S. cerevisiae SET3 complex includes two histone deacetylases, Hos2 and Hst1, and is a meiotic-specific repressor of the sporulation gene program. Genes Dev 2001, 15:2991-3004.

32. Rincon-Arano H, Halow J, Delrow JJ, Parkhurst SM, Groudine M: UpSET recruits HDAC complexes and restricts chromatin accessibility and acetylation at promoter regions. Cell 2012, 151:1214–1228.

33. McElroy KA, Jung YL, Zee BM, Wang CI, Park PJ, Kuroda MI: upSET, the Drosophila homologue of SET3, Is Required for Viability and the Proper Balance of Active and Repressive Chromatin Marks. G3 (Bethesda) 2017, 7:625-635.

34. Ni Z, Ebata A, Alipanahiramandi E, Lee SS: Two SET domain containing genes link epigenetic changes and aging in Caenorhabditis elegans. Aging Cell 2012, 11:315–325.

35. Emerson FJ, Chiu C, Lin LY, Riedel CG, Zhu M, Lee SS: The chromatin factors SET-26 and HCF-1 oppose the histone deacetylase HDA-1 in longevity and gene regulation in C. elegans. bioRxiv 2023.

36. Tsur A, Bening Abu-Shach U, Broday L: ULP-2 SUMO Protease Regulates E-Cadherin Recruitment to Adherens Junctions. Developmental Cell 2015, 35:63–77.

37. Wagner K, Kunz K, Piller T, Tascher G, Hölper S, Stehmeier P, Keiten-Schmitz J, Schick M, Keller U, Müller S: The SUMO Isopeptidase SENP6 Functions as a Rheostat of Chromatin Residency in Genome Maintenance and Chromosome Dynamics. Cell Rep 2019, 29:480–494.e485.

38. Mitra S, Bodor DL, David AF, Abdul-Zani I, Mata JF, Neumann B, Reither S, Tischer C, Jansen LET: Genetic screening identifies a SUMO protease dynamically maintaining centromeric chromatin. Nat Commun 2020, 11:501.

39. Liebelt F, Jansen NS, Kumar S, Gracheva E, Claessens LA, Verlaan-de Vries M, Willemstein E, Vertegaal ACO: The poly-SUMO2/3 protease SENP6 enables assembly of the constitutive centromere-associated network by group deSUMOylation. Nat Commun 2019, 10:3987.

40. Hsin H, Kenyon C: Signals from the reproductive system regulate the lifespan of C. elegans. Nature 1999, 399:362–366.

41. Antebi A: Regulation of longevity by the reproductive system. Exp Gerontol 2013, 48:596-602.

42. Knutson AK, Egelhofer T, Rechtsteiner A, Strome S: Germ Granules Prevent Accumulation of Somatic Transcripts in the Adult Caenorhabditis elegans Germline. Genetics 2017, 206:163–178.

43. Iwasaki K, McCarter J, Francis R, Schedl T: emo-1, a Caenorhabditis elegans Sec61p gamma homologue, is required for oocyte development and ovulation. J Cell Biol 1996, 134:699–714.

44. Tocchini C, Keusch JJ, Miller SB, Finger S, Gut H, Stadler MB, Ciosk R: The TRIM-NHL protein LIN-41 controls the onset of developmental plasticity in Caenorhabditis elegans. PLoS Genet 2014, 10:e1004533.

45. Maduro M, Pilgrim D: Identification and cloning of unc-119, a gene expressed in the Caenorhabditis elegans nervous system. Genetics 1995, 141:977–988.

46. Updike Dustin L, Knutson Andrew Ka, Egelhofer Thea A, Campbell Anne C, Strome S: Germ-Granule Components Prevent Somatic Development in the *C.&#xa0;elegans* Germline. Current Biology 2014, 24:970–975.

47. Ciosk R, DePalma M, Priess JR: Translational Regulators Maintain Totipotency in the Caenorhabditis elegans Germline. Science 2006, 311:851–853.

48. Tursun B, Patel T, Kratsios P, Hobert O: Direct Conversion of C. elegans Germ Cells into Specific Neuron Types. Science 2011, 331:304-308.

49. Ul Fatima N, Tursun B: Conversion of Germ Cells to Somatic Cell Types in C. elegans. J Dev Biol 2020, 8.

50. Santos-Rosa H, Schneider R, Bannister AJ, Sherriff J, Bernstein BE, Emre NC, Schreiber SL, Mellor J, Kouzarides T: Active genes are tri-methylated at K4 of histone H3. Nature 2002, 419:407–411.

51. Wiles ET, Selker EU: H3K27 methylation: a promiscuous repressive chromatin mark. Curr Opin Genet Dev 2017, 43:31–37.

52. Patel T, Tursun B, Rahe DP, Hobert O: Removal of Polycomb repressive complex 2 makes C. elegans germ cells susceptible to direct conversion into specific somatic cell types. Cell Rep 2012, 2:1178–1186.

53. Hickey CM, Wilson NR, Hochstrasser M: Function and regulation of SUMO proteases. Nat Rev Mol Cell Biol 2012, 13:755–766.

54. Jumper J, Evans R, Pritzel A, Green T, Figurnov M, Ronneberger O, Tunyasuvunakool K, Bates R, Žídek A, Potapenko A, Bridgland A, Meyer C, Kohl SAA, Ballard AJ, Cowie A, Romera-Paredes B, Nikolov S, Jain R, Adler J, Back T, Petersen S, Reiman D, Clancy E, Zielinski M, Steinegger M, Pacholska M, Berghammer T, Bodenstein S, Silver D, Vinyals O, Senior AW, Kavukcuoglu K, Kohli P, Hassabis D: Highly accurate protein structure prediction with AlphaFold. Nature 2021, 596:583–589.

55. Hendriks IA, Lyon D, Young C, Jensen LJ, Vertegaal AC, Nielsen ML: Site-specific mapping of the human SUMO proteome reveals co-modification with phosphorylation. Nat Struct Mol Biol 2017, 24:325–336.

56. Andersen EC, Horvitz HR: Two C. elegans histone methyltransferases repress lin-3 EGF transcription to inhibit vulval development. Development 2007, 134:2991–2999.

57. Reichman R, Shi Z, Malone R, Smolikove S: Mitotic and Meiotic Functions for the SUMOylation Pathway in the Caenorhabditis elegans Germline. Genetics 2018, 208:1421–1441.

58. Michaeli L, Spector E, Haeussler S, Carvalho CA, Grobe H, Abu-Shach UB, Zinger H, Conradt B, Broday L: ULP-2 SUMO protease regulates UPR(mt) and mitochondrial homeostasis in Caenorhabditis elegans. Free Radic Biol Med 2024, 214:19–27.

59. Capowski EE, Martin P, Garvin C, Strome S: Identification of grandchildless loci whose products are required for normal germ-line development in the nematode Caenorhabditis elegans. Genetics 1991, 129:1061–1072.

60. Bender LB, Cao R, Zhang Y, Strome S: The MES-2/MES-3/MES-6 complex and regulation of histone H3 methylation in C. elegans. Curr Biol 2004, 14:1639–1643.

61. Zhang X, Novera W, Zhang Y, Deng LW: MLL5 (KMT2E): structure, function, and clinical relevance. Cell Mol Life Sci 2017, 74:2333–2344.

62. Katz DJ, Edwards TM, Reinke V, Kelly WG: A C. elegans LSD1 demethylase contributes to germline immortality by reprogramming epigenetic memory. Cell 2009, 137:308–320.

63. Alvares SM, Mayberry GA, Joyner EY, Lakowski B, Ahmed S: H3K4 demethylase activities repress proliferative and postmitotic aging. Aging Cell 2014, 13:245–253.

64. Robert VJ, Mercier MG, Bedet C, Janczarski S, Merlet J, Garvis S, Ciosk R, Palladino F: The SET-2/SET1 histone H3K4 methyltransferase maintains pluripotency in the Caenorhabditis elegans germline. Cell Rep 2014, 9:443–450.

65. Saltzman AL, Soo MW, Aram R, Lee JT: Multiple Histone Methyl-Lysine Readers Ensure Robust Development and Germline Immortality in Caenorhabditis elegans. Genetics 2018, 210:907–923.

66. Kim H, Ding YH, Lu S, Zuo MQ, Tan W, Conte D, Jr., Dong MQ, Mello CC: PIE-1 SUMOylation promotes germline fates and piRNA-dependent silencing in C. elegans. Elife 2021, 10.

67. Kim H, Ding YH, Zhang G, Yan YH, Conte D, Jr., Dong MQ, Mello CC: HDAC1 SUMOylation promotes Argonaute-directed transcriptional silencing in C. elegans. Elife 2021, 10.

68. Jambhekar A, Dhall A, Shi Y: Roles and regulation of histone methylation in animal development. Nat Rev Mol Cell Biol 2019, 20:625–641.

69. Peng J, Wysocka J: It takes a PHD to SUMO. Trends in Biochemical Sciences 2008, 33:191–194.

70. Miura K, Renhu N, Suzaki T: The PHD finger of Arabidopsis SIZ1 recognizes trimethylated histone H3K4 mediating SIZ1 function and abiotic stress response. Commun Biol 2020, 3:23.

71. Ryu HY, Hochstrasser M: Histone sumoylation and chromatin dynamics. Nucleic Acids Res 2021, 49:6043–6052.

72. Ryu HY, Zhao D, Li J, Su D, Hochstrasser M: Histone sumoylation promotes Set3 histone-deacetylase complex-mediated transcriptional regulation. Nucleic Acids Res 2020, 48:12151–12168.

73. Nayak A, Viale-Bouroncle S, Morsczeck C, Muller S: The SUMO-specific isopeptidase SENP3 regulates MLL1/MLL2 methyltransferase complexes and controls osteogenic differentiation. Mol Cell 2014, 55:47–58.

74. Nayak A, Lopez-Davila AJ, Kefalakes E, Holler T, Kraft T, Amrute-Nayak M: Regulation of SETD7 Methyltransferase by SENP3 Is Crucial for Sarcomere Organization and Cachexia. Cell Rep 2019, 27:2725–2736.e2724.

75. Brenner S: The genetics of Caenorhabditis elegans. Genetics 1974, 77:71-94.

76. Dejima K, Hori S, Iwata S, Suehiro Y, Yoshina S, Motohashi T, Mitani S: An Aneuploidy-Free and Structurally Defined Balancer Chromosome Toolkit for Caenorhabditis elegans. Cell Rep 2018, 22:232–241.

77. Savion N, Levine A, Kotev-Emeth S, Bening Abu-Shach U, Broday L: S-allylmercapto-N-acetylcysteine protects against oxidative stress and extends lifespan in Caenorhabditis elegans. PLoS One 2018, 13:e0194780.

78. Pelisch F, Hay RT: Tools to Study SUMO Conjugation in Caenorhabditis elegans. Methods Mol Biol 2016, 1475:233–256.

79. Vidal M, Brachmann RK, Fattaey A, Harlow E, Boeke JD: Reverse two-hybrid and one-hybrid systems to detect dissociation of protein-protein and DNA-protein interactions. Proc Natl Acad Sci U S A 1996, 93:10315–10320.

80. Yu H, Braun P, Yildirim MA, Lemmens I, Venkatesan K, Sahalie J, Hirozane-Kishikawa T, Gebreab F, Li N, Simonis N, Hao T, Rual JF, Dricot A, Vazquez A, Murray RR, Simon C, Tardivo L, Tam S, Svrzikapa N, Fan C, de Smet AS, Motyl A, Hudson ME, Park J, Xin X, Cusick ME, Moore T, Boone C, Snyder M, Roth FP, Barabási AL, Tavernier J, Hill DE, Vidal M: High-quality binary protein interaction map of the yeast interactome network. Science 2008, 322:104–110.

81. Vidalain PO, Boxem M, Ge H, Li S, Vidal M: Increasing specificity in high-throughput yeast two-hybrid experiments. Methods 2004, 32:363–370.

82. Koorman T, Klompstra D, van der Voet M, Lemmens I, Ramalho JJ, Nieuwenhuize S, van den Heuvel S, Tavernier J, Nance J, Boxem M: A combined binary interaction and phenotypic map of C. elegans cell polarity proteins. Nat Cell Biol 2016, 18:337–346.

83. Reverter D, Lima CD: Preparation of SUMO proteases and kinetic analysis using endogenous substrates. Methods Mol Biol 2009, 497:225–239.

84. Paix A, Folkmann A, Seydoux G: Precision genome editing using CRISPR-Cas9 and linear repair templates in C. elegans. Methods 2017, 121–122:86-93.

